# Adult Stem Cell-derived Complete Lung Organoid Models Emulate Lung Disease in COVID-19

**DOI:** 10.1101/2020.10.17.344002

**Authors:** Courtney Tindle, MacKenzie Fuller, Ayden Fonseca, Sahar Taheri, Stella-Rita Ibeawuchi, Nathan Beutler, Gajanan D. Katkar, Amanraj Claire, Vanessa Castillo, Moises Hernandez, Hana Russo, Jason Duran, Laura E. Crotty Alexander, Ann Tipps, Grace Lin, Patricia A. Thistlethwaite, Ranajoy Chattopadhyay, Thomas F. Rogers, Debashis Sahoo, Pradipta Ghosh, Soumita Das

## Abstract

SARS-CoV-2, the virus responsible for COVID-19, causes widespread damage in the lungs in the setting of an overzealous immune response whose origin remains unclear. We present a scalable, propagable, personalized, cost-effective adult stem cell-derived human lung organoid model that is complete with both proximal and distal airway epithelia. Monolayers derived from adult lung organoids (ALOs), primary airway cells, or hiPSC-derived alveolar type-II (AT2) pneumocytes were infected with SARS-CoV-2 to create *in vitro* lung models of COVID-19. Infected ALO-monolayers best recapitulated the transcriptomic signatures in diverse cohorts of COVID-19 patient-derived respiratory samples. The airway (proximal) cells were critical for sustained viral infection, whereas distal alveolar differentiation (AT2→AT1) was critical for mounting the overzealous host immune response in fatal disease; ALO monolayers with well-mixed proximodistal airway components recapitulated both. Findings validate a human lung model of COVID-19, which can be immediately utilized to investigate COVID-19 pathogenesis and vet new therapies and vaccines.

**GRAPHIC ABSTRACT:** 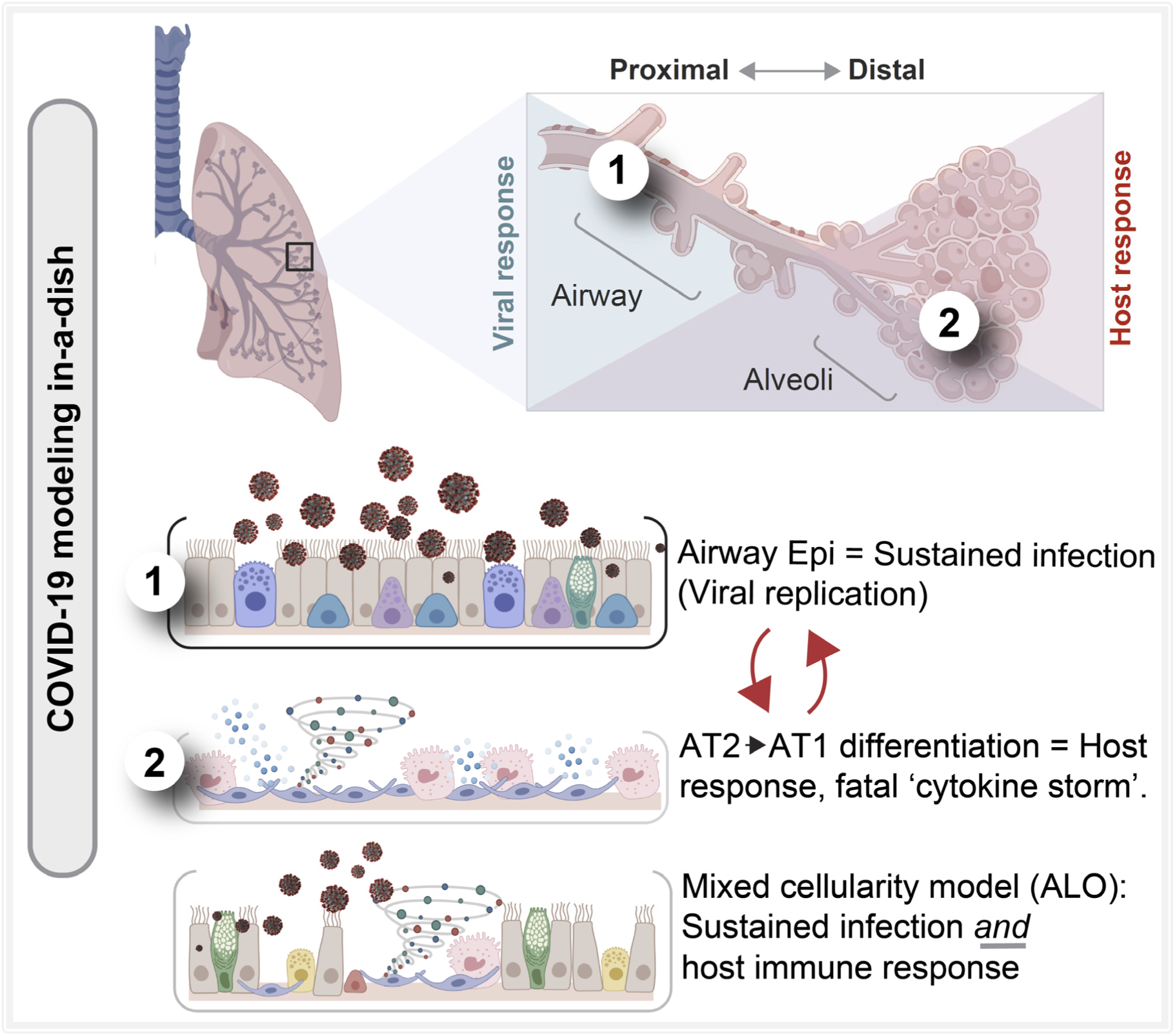

**HIGHLIGHTS:**

- Human lung organoids with mixed proximodistal epithelia are created
- Proximal airway cells are critical for viral infectivity
- Distal alveolar cells are important for emulating host response
- Both are required for the overzealous response in severe COVID-19

**IN BRIEF:** An integrated stem cell-based disease modeling and computational approach demonstrate how both proximal airway epithelium is critical for SARS-CoV-2 infectivity, but distal differentiation of alveolar pneumocytes is critical for simulating the overzealous host response in fatal COVID-19.

## Introduction

SARS-CoV-2, the virus responsible for COVID-19, causes widespread inflammation and injury in the lungs, giving rise to diffuse alveolar damage (DAD) ^1–5^, featuring marked infection and viral burden leading to apoptosis of alveolar pneumocytes ^6^, along with pulmonary edema ^7, 8^. DAD leads to poor gas exchange and, ultimately, respiratory failure; the latter appears to be the final common mechanism of death in most patients with severe COVID-19 infection. How the virus causes so much damage remains unclear. A particular challenge is to understand the out-of-control immune reaction to the SARS-CoV-2 infection known as a cytokine storm, which has been implicated in many of the deaths from COVID-19. Although rapidly developed pre-clinical animal models have recapitulated some of the pathognomonic aspects of infection, e.g., induction of disease, and transmission, and even viral shedding in the upper and lower respiratory tract, many failed to develop severe clinical symptoms ^9^. Thus, the need for pre-clinical models remains both urgent and unmet.

To address this need, several groups have attempted to develop human pre-clinical COVID-19 lung models, all within the last few months ^10–12^. While a head-to-head comparison of the key characteristics of each model can be found in **Table 1**, what is particularly noteworthy is that none recapitulate the heterogeneous epithelial cellularity of both proximal and distal airways, i.e., airway epithelia, basal cells, secretory club cells and alveolar pneumocytes. Also noteworthy is that models derived from iPSCs lack propagability and/or cannot be reproducibly generated for biobanking; nor can they be scaled up in cost-effective ways for use in drug screens. Besides the approaches described so far, there are a few more approaches used for modeling COVID-19—(i) 3D organoids from bronchospheres and tracheospheres have been established before ^13–15^ and are now used in apical-out cultures for infection with SARS-COV-2 ^16^; (ii) the most common model used for drug screening is the air-liquid interphase (ALI model) in which pseudo-stratified primary bronchial or small airway epithelial cells are used to recreate the multilayered mucociliary epithelium ^17, 18^; (iii) several groups have also generated 3D airway models from iPSCs or tissue-resident stem cells ^19–24^; (iv) others have generated AT2 cells from iPSCs using closely overlapping protocols of sequential differentiation starting with definitive endoderm, anterior foregut endoderm, and distal alveolar expression ^25–30^. (v) Finally, long term *in vitro* culture conditions for pseudo-stratified airway epithelium organoids, derived from healthy and diseased adult humans suitable to assess virus infectivity ^31–33^ have been pioneered; unfortunately, these airway organoids expressed virtually no lung mesenchyme or alveolar signature. What remains unclear is if any of these models accurately recapitulate the immunopathologic phenotype that is seen in the lungs in COVID-19.

**Table 1.**
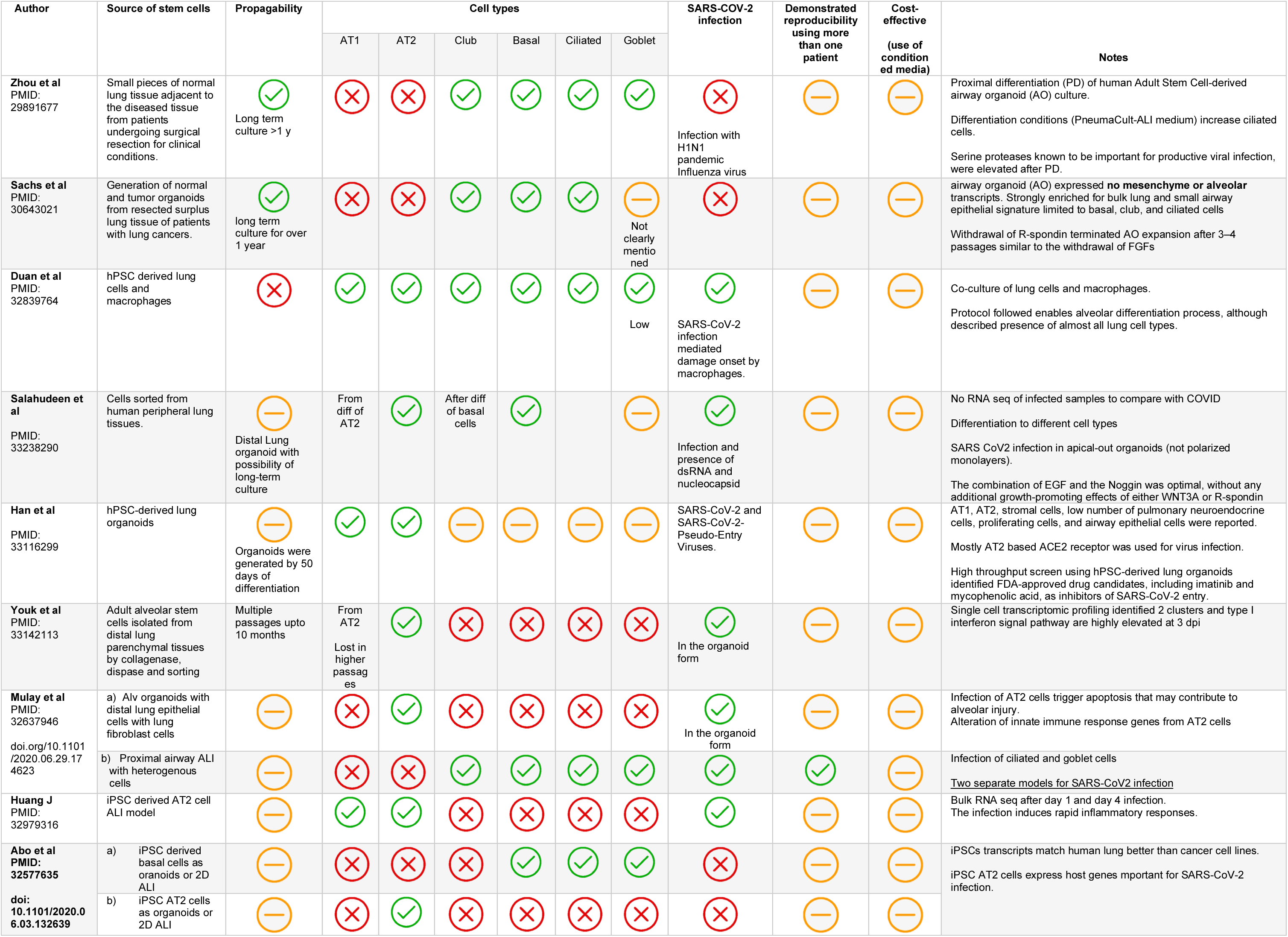
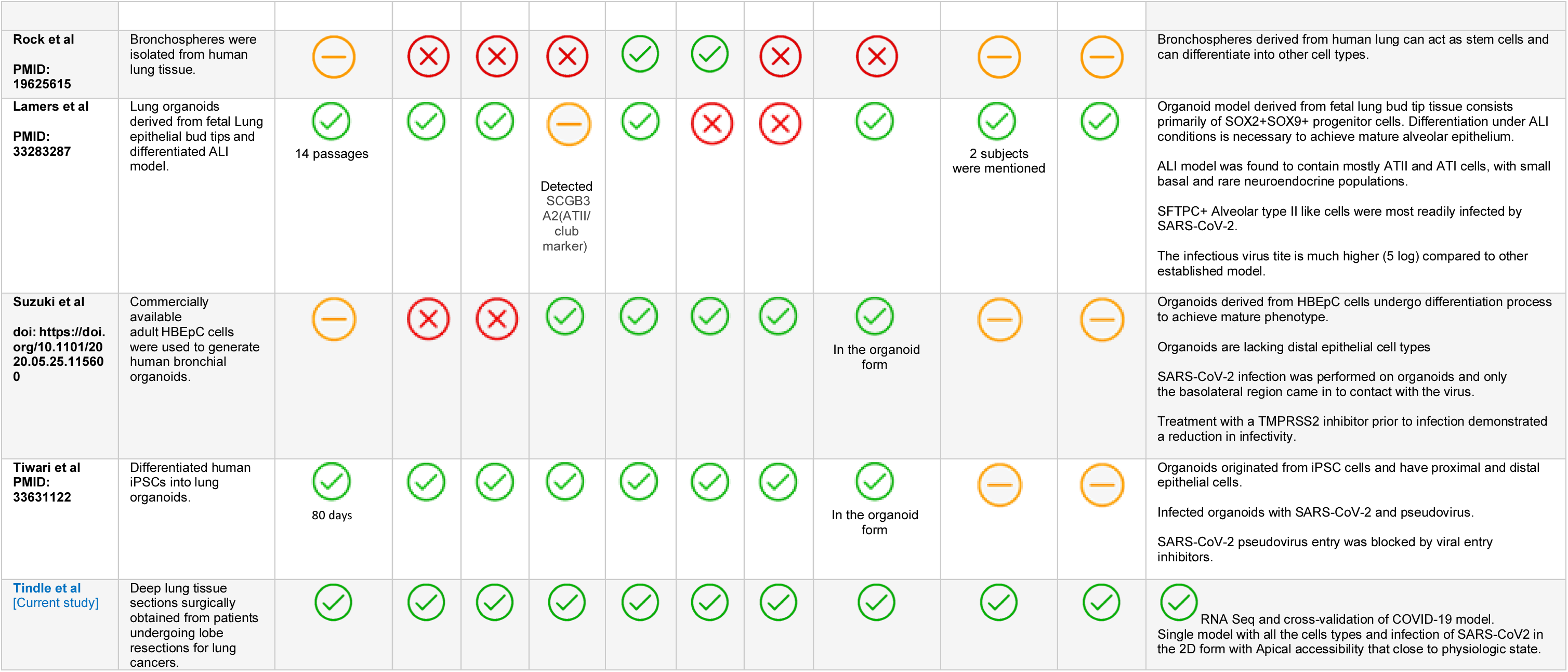
A comparison of current versus existing lung organoid models available for modeling COVID-19

We present a rigorous transdisciplinary approach that systematically assesses an adult lung organoid model that is propagable, personalized and complete with both proximal airway and distal alveolar cell types against existing models that are incomplete, and we cross-validate them all against COVID-19 patient-derived respiratory samples. Findings surprisingly show that cellular crosstalk between both proximal and distal components are necessary to emulate how SARS-CoV-2 causes diffuse alveolar pneumocyte damage; the proximal airway mounts a sustained viral infection, but it is the distal alveolar pneumocytes that mount the overzealous host response that has been implicated in a fatal disease.

## Results

### A rationalized approach for creating and validating acute lung injury in COVID-19

To determine which cell types in the lungs might be most readily infected, we began by analyzing a human lung single-cell sequencing dataset (GSE132914) for the levels of expression of angiotensin-converting enzyme-II (ACE2) and Transmembrane Serine Protease 2 (TMPRSS2), the two receptors that have been shown to be the primary sites of entry for the SARS-CoV-2 ^34^. The dataset was queried with widely accepted markers of all the major cell types (see **Table 2**). Alveolar epithelial type 2 (AT2), ciliated and club cells emerged as the cells with the highest expression of both receptors (**Fig. 1A; Figure 1- figure supplement 1A**). These observations are consistent with published studies demonstrating that ACE2 is indeed expressed highest in AT2 and ciliated cells ^11, 35, 36^. In a cohort of deceased COVID-19 patients, we observed by H&E (**Figure 1- figure supplement 1B**) that gas-exchanging flattened AT1 pneumocytes are virtually replaced by cuboidal cells that were subsequently confirmed to be AT2-like cells via immunofluorescent staining with the AT2-specific marker, surfactant protein-C (SFTPC; **Fig. 1B** *upper panel***; Figure 1- figure supplement 1C**; *top*). We also confirmed that club cells express ACE2 (**Figure 1- figure supplement 1C**; *bottom*), underscoring the importance of preserving these cells in any ideal lung model of COVID-19. When we analyzed the lungs of deceased COVID-19 patients, the presence of SARS-COV-2 in alveolar pneumocytes was also confirmed, as determined by the colocalization of viral nucleocapsid protein with SFTPC (**Fig. 1B**; *lower panel;* **Figure 1- figure supplement 1D**). Immunohistochemistry studies further showed the presence of SARS-COV-2 virus in alveolar pneumocytes and in alveolar immune cells (**Figure 1- figure supplement 1E**). These findings are consistent with the gathering consensus that alveolar pneumocytes support the interaction between the epithelial cells and inflammatory cells recruited to the lung; via mechanisms that remain unclear, they are generally believed to contribute to the development of acute lung injury and acute respiratory distress syndrome (ARDS), the severe hypoxemic respiratory failure during COVID-19 ^37, 38^. Because prior work has demonstrated that SARS-CoV-2 infectivity in patient-derived airway cells is highest in the proximal airway epithelium compared to the distal alveolar pneumocytes (AT1 and AT2) ^37^, and yet, it is the AT2 pneumocytes that harbor the virus, and the AT1 pneumocytes that are ultimately destroyed during diffuse alveolar damage, we hypothesized that both proximal airway and distal (alveolar pneumocyte) components might play distinct roles in the respiratory system to mount the so-called viral infectivity and host immune response phases of the clinical symptoms observed in COVID-19^39^.

**Figure 1.**
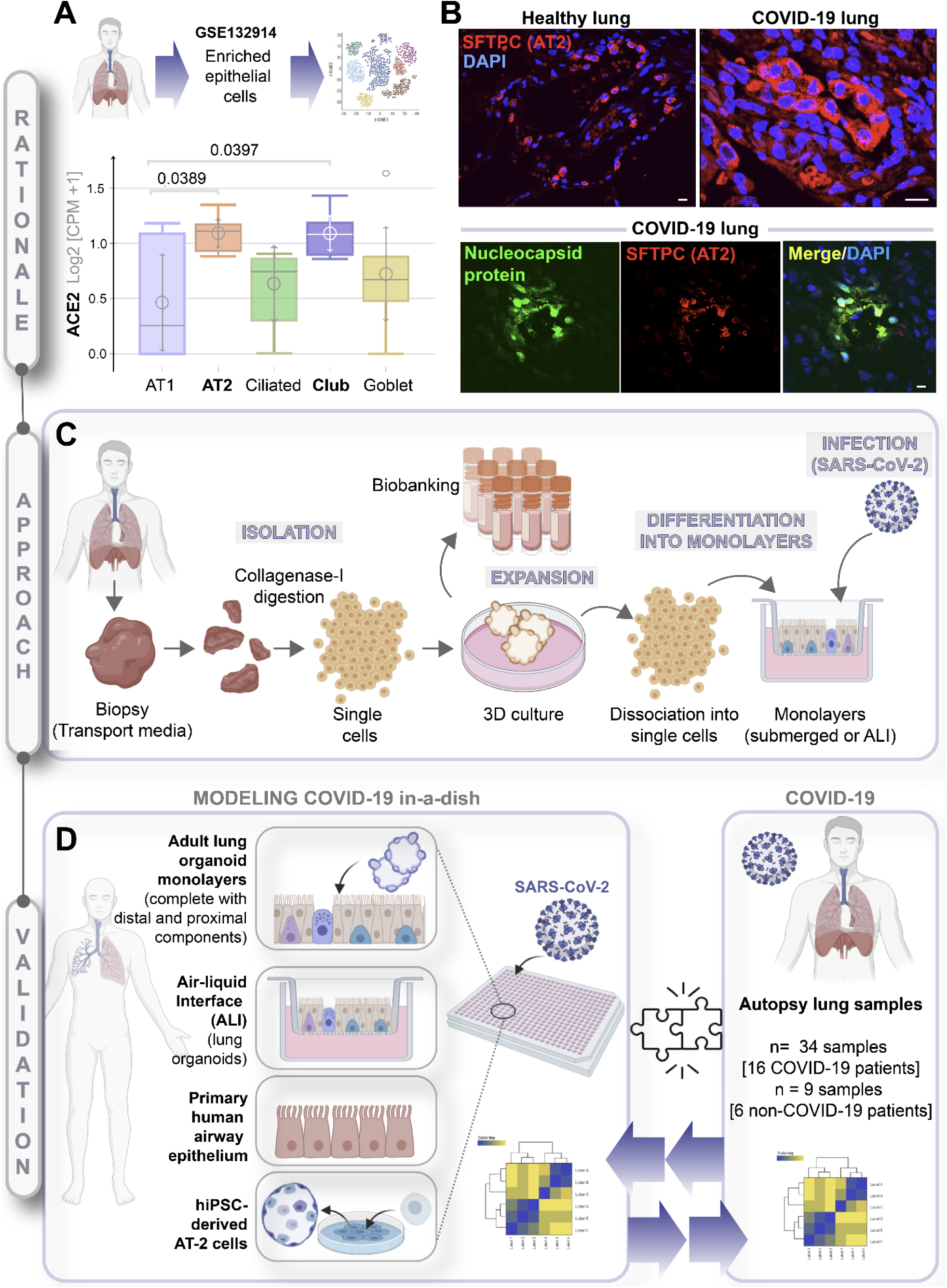
**A rationalized approach to building and validating human pre-clinical models of COVID-19. A**. Whisker plots display relative levels of ACE2 expression in various cell types in the normal human lung. The cell types were annotated within a publicly available single-cell sequencing dataset (GSE132914) using genes listed in **Table 1**. *p* values were analyzed by one-way Anova and Tukey post hoc test. **B.** FFPE sections of the human lung from normal and deceased COVID-19 patients were stained for SFTPC, alone or in combination with nucleocapsid protein and analyzed by confocal immunofluorescence. Representative images are shown. Scale bar = 20 µm. **C.** Schematic showing key steps generating an adult stem cell-derived, propagable, lung organoid model, complete with proximal and distal airway components for modeling COVID-19-in-a-dish. See *Methods* for details regarding culture conditions. **D.** A transcriptome-based approach is used for cross-validation of *in vitro* lung models of SARS-CoV-2 infection (left) versus the human disease, COVID-19 (right), looking for a match in gene expression signatures.

**Table 2:**
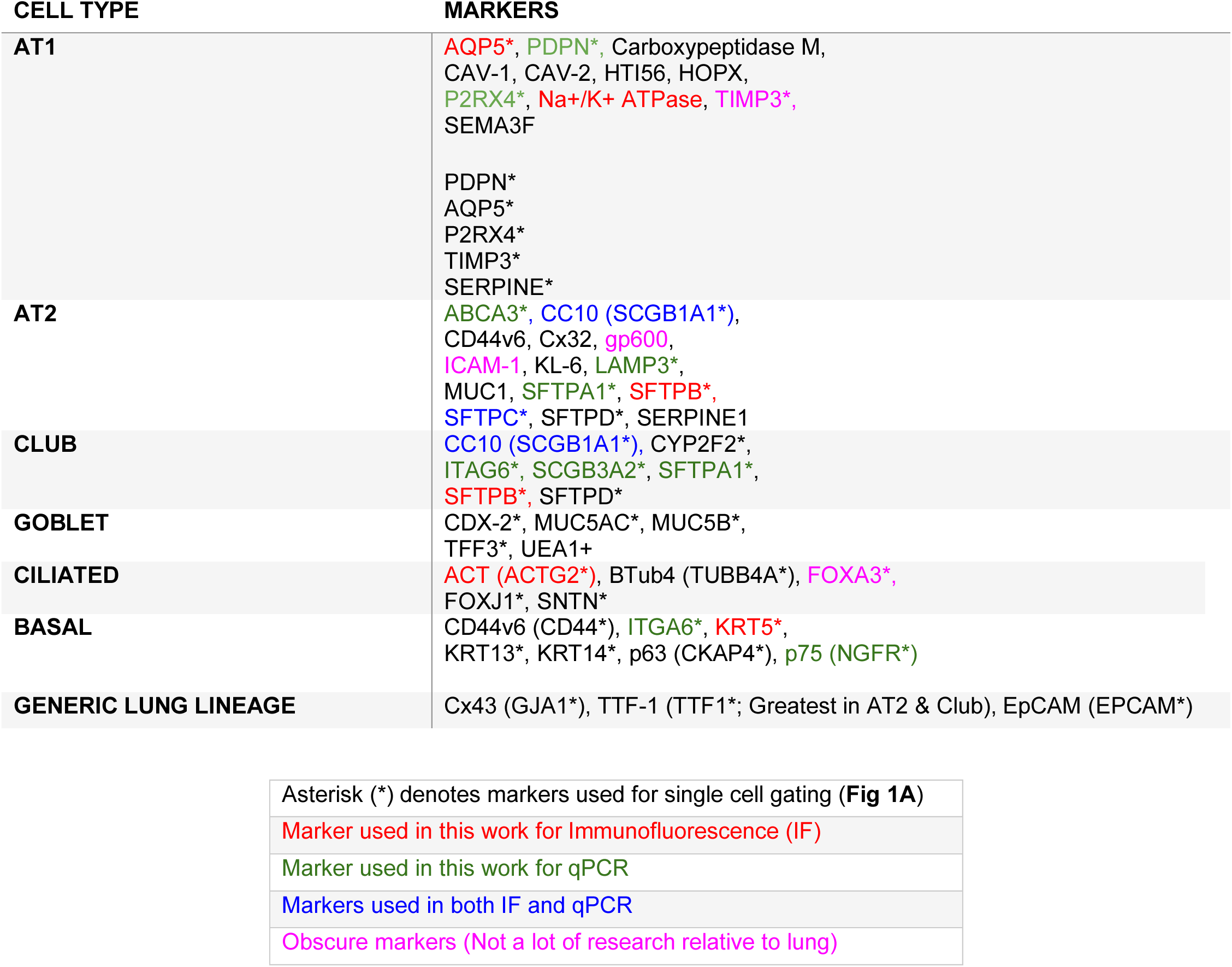
Markers used to identify various cell types in the lung

| CELL TYPE | MARKERS |
| --- | --- |
| AT1 | AQP5*, PDPN*, Carboxypeptidase M, CAV-1, CAV-2, HTI56, HOPX, P2RX4*, Na+/K+ ATPase, TIMP3*, SEMA3F<br>PDPN*<br>AQP5*<br>P2RX4*<br>TIMP3*<br>SERPINE* |
| AT2 | ABCA3*, CC10 (SCGB1A1*), CD44v6, Cx32, gp600, ICAM-1, KL-6, LAMP3*, MUC1, SFTPA1*, SFTPB*, SFTPC*, SFTPD*, SERPINE1 |
| CLUB | CC10 (SCGB1A1*), CYP2F2*, ITAG6*, SCGB3A2*, SFTPA1*, SFTPB*, SFTPD* |
| GOBLET | CDX-2*, MUC5AC*, MUC5B*, TFF3*, UEA1+ |
| CILIATED | ACT (ACTG2*), BTub4 (TUBB4A*), FOXA3*, FOXJ1*, SNTN* |
| BASAL | CD44v6 (CD44*), ITGA6*, KRT5*, KRT13*, KRT14*, p63 (CKAP4*), p75 (NGFR*) |
| GENERIC LUNG LINEAGE | Cx43 (GJA1*), TTF-1 (TTF1*; Greatest in AT2 & Club), EpCAM (EPCAM*) |

|  |
| --- |
| Asterisk (*) denotes markers used for single cell gating (**Fig 1A**) |
| Marker used in this work for Immunofluorescence (IF) |
| Marker used in this work for qPCR |
| Markers used in both IF and qPCR |
| Obscure markers (Not a lot of research relative to lung) |

Because no existing lung model provides such proximodistal cellular representation (**Table 1**), and hence, may not recapitulate with accuracy the clinical phases of COVID-19, we first sought to develop a lung model that is complete with both proximal and distal airway epithelia using adult stem cells that were isolated from deep lung biopsies (i.e., sufficient to reach the bronchial tree). Lung organoids were generated using the workflow outlined in **Fig 1C** and a detailed protocol that had key modifications from previously published ^31, 33^ methodologies (see *Methods*). Organoids grown in 3D cultures were subsequently dissociated into single cells to create 2D-monolayers (either maintained submerged in media or used in ALI model) for SARS-CoV-2 infection, followed by RNA Seq analysis. Primary airway epithelial cells and hiPSC-derived alveolar type-II (AT2) pneumocytes were used as additional models (**Fig. 1D**; *left panel*). Each of these transcriptomic datasets was subsequently used to cross-validate our *ex-vivo* lung models of SARS-CoV-2 infection with the human COVID-19 autopsy lung specimens (**Fig. 1D**; *right panel*) to objectively vet each model for their ability to accurately recapitulate the gene expression signatures in the patient-derived lungs.

### Creation of a lung organoid model, complete with both proximal and distal airway epithelia

Three lung organoid lines were developed from deep lung biopsies obtained from the normal regions of lung lobes surgically resected for lung cancer; both genders, smokers and non-smokers were represented (**Figure 2- figure supplement1A; Table 3**). Three different types of media were compared (**Figure 2- figure supplement 1B**); the composition of these media was inspired either by their ability to support adult-stem cell-derived mixed epithelial cellularity in other organs (like the gastrointestinal tract ^40–42^, or rationalized based on published growth conditions for proximal and distal airway components ^25, 31, 32^. A growth condition that included conditioned media from L-WRN cells which express Wnt3, R-spondin and Noggin, supplemented with recombinant growth factors, which we named as ‘*lung organoid expansion media*’ emerged as superior compared to alveolosphere media-I and II^27, 28^ (details in the methods), based on its ability to consistently and reproducibly support the best morphology and growth characteristics across multiple attempts to isolate organoids from lung tissue samples. Three adult lung organoid lines (ALO1-3) were developed using the expansion media, monitored for their growth characteristics by brightfield microscopy and cultured with similar phenotypes until P10 and beyond (**Figure 2- figure supplement 1C-D**). The 3D morphology of the lung organoid was also assessed by H&E staining of slices cut from formalin-fixed paraffin-embedded (FFPE) cell blocks of *HistoGel*-embedded ALO1-3 (**Figure 2- figure supplement 1E**).

**Table 3.** Characteristics of patients enrolled into this study for obtaining lung tissues to serve as source of stem cells to generate lung organoids

| **<u>Name</u>** | **<u>Date of surgery</u>** | **<u>Age</u>** | **<u>Sex</u>** | **<u>Smoking History</u>** | **<u>Reason for surgery</u>** | **<u>Histology</u>** |
| --- | --- | --- | --- | --- | --- | --- |
| **ALO1** | 4/17/2020 | 64 | Male | Current, chronic smoker.<br>Packs/day: 0.50.<br>Years: 53.<br>Pack years: 26.5 | Lung Carcinoma | Invasive Squamous Cell Carcinoma, non-keratinizing |
| **ALO2** | 4/17/2020 | 59 | Male | Non-smoker | Lung Carcinoma | Invasive Adenocarcinoma |
| **ALO3** | 7/7/2020 | 46 | Female | Non-smoker | Left Lower Lobe Nodule | Invasive Adenocarcinoma |

To determine if all the 6 major lung epithelial cells (illustrated in **Fig. 2A**) are present in the organoids, we analyzed various cell-type markers by qRT-PCR (**Fig. 2B-H; Figure 2- figure supplement 2A-H**). All three ALO lines had a comparable level of AT2 cell surfactant markers (compared against hiPSC-derived AT2 cells as positive control) and a significant amount of AT1, as determined using the marker AQP5. ALOs also contained basal cells (as determined by the marker ITGA6 and p75/NGFR), ciliated cells (as determined by the marker FOXJ1), club cells (as determined by the marker SCGB1A1) and stem cells (as determined by marker TP63). As expected, the primary human bronchial epithelial cells (NHBE) had significantly higher expression of basal cell markers than the ALO lines (hence, served as a positive control), but they lacked stemness and club cells (hence, served as a negative control).

**Figure 2.**
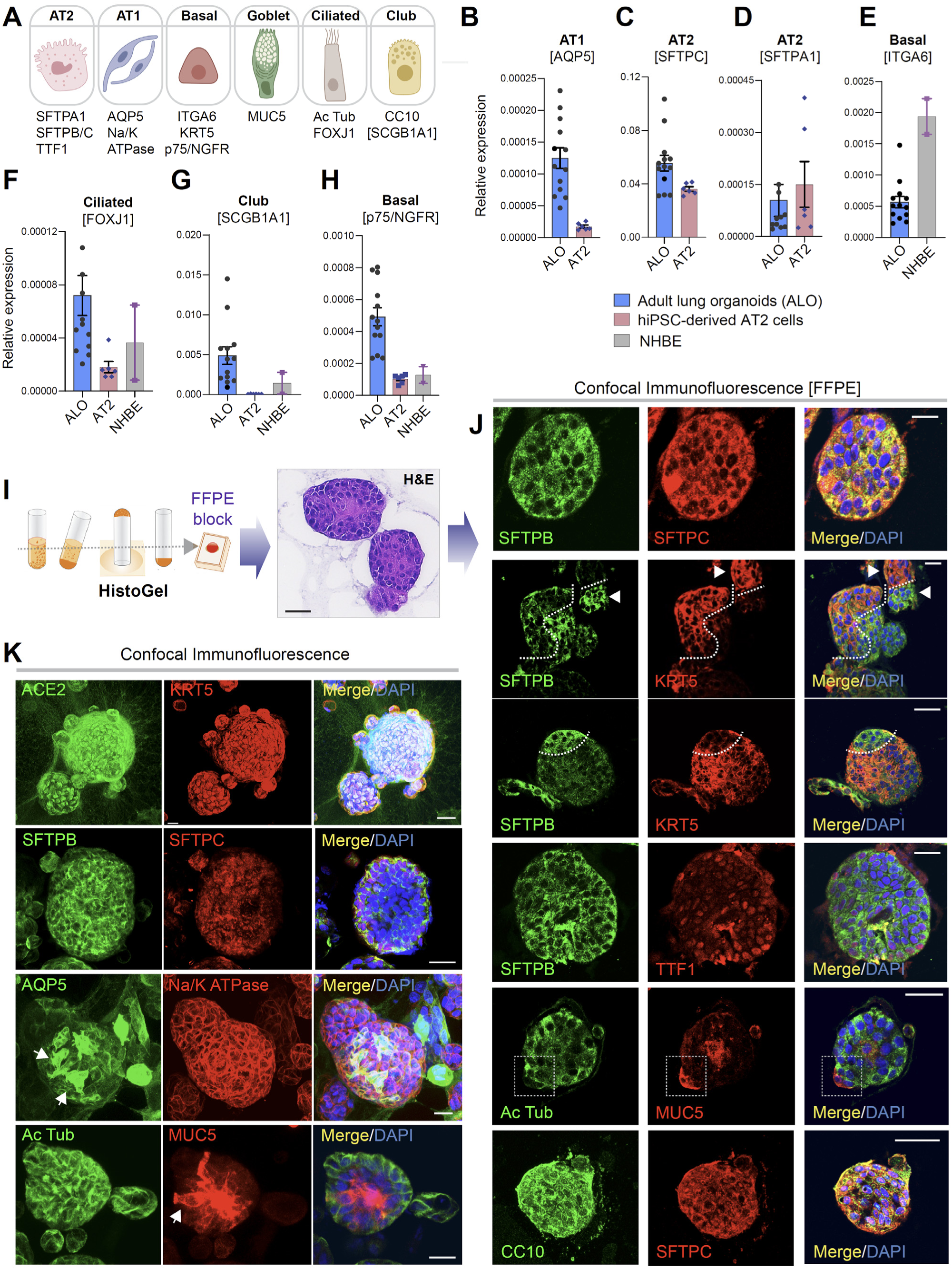
**Adult stem cell-derived lung organoids are propagatable models with both proximal and distal airway components**. **A.** Schematic lists the various markers used here for qPCR and immunofluorescence to confirm the presence of all cell types in the 3D lung organoids here and in 2D monolayers later (in Fig 3). **B-H.** Bar graphs display the relative abundance of various cell type markers (normalized to 18S) in adult lung organoids (ALO), compared to the airway (NHBE) and/or alveolar (AT2) control cells, as appropriate. *p* values were analyzed by one-way Anova. Error bars denote S.E.M; n = 3-6 datasets from 3 independent ALOs and representing early and late passages. See also **Figure 2- figure supplement 2** for individual ALOs. **I-J**. H&E*-* stained cell blocks were prepared using *HistoGel* (I). Slides were stained for the indicated markers and visualized by confocal immunofluorescence microscopy. Representative images are shown in J. Scale bar = 50 µm. **K**. 3D organoids grown in 8-well chamber slides were fixed, immunostained and visualized by confocal microscopy as in J. Scale bar = 50 µm. See also **Figure 2- figure supplement 2**. Single channel images are max projections of z-stacks; however, merged panels are optical sections to visualize the centers of the organoids. All immunofluorescence images showcased in this figure were obtained from ALO lines within passage #3-6. See also **Figure 2- figure supplement 3-5** for additional evidence of mixed cellularity of ALO models, their similarity to lung tissue of origin, and stability of cellular composition during early (#1-8) and late (#8-15) passages, as determined by qPCR and flow cytometry.

The presence of all cell types was also confirmed by assessing protein expression of various cell types within organoids grown in 3D cultures. Two different approaches were used—(i) slices cut from FFPE cell blocks of *HistoGel*-embedded ALO lines (**Fig. 2I-J**) or (ii) ALO lines grown in 8-well chamber slides were fixed in matrigel (**Fig. 2K**), stained, and assessed by confocal microscopy. Such staining not only confirmed the presence of all cell types in each ALO line but also demonstrated the presence of more than one cell type (i.e., mixed cellularity) of proximal (basal-KRT5) and distal (AT1/AT2 markers) within the same organoid structure. For example, AT2 and basal cells, marked by SFTPB and KRT5, respectively, were found in the same 3D structure (**Fig. 2J**, interrupted curved lines). Similarly, ciliated cells and goblet cells stained by Ac-Tub and Muc5, respectively, were found to coexist within the same structure (**Fig. 2J**, interrupted box; **Fig. 2K**, arrow). Intriguingly, we noted 3D structures comprised of cells that co-stained for CC10 and SFTPC (**Fig 2J**, bottom panel), likely representative of a unique population of multipotent stem cells termed bronchioalveolar stem cells (BASCs), which have been found to be located at the bronchioalveolar-duct junctions (BADJs)^43, 44^. Besides the organoids with heterogeneous makeup, each ALO line also showed homotypic organoid structures that were relatively enriched in one cell type (**Fig. 2J**, arrowheads pointing to two adjacent structures that are either KRT5- or SFTPB- positive). Regardless of their homotypic or heterotypic cellular organization into 3D structures, the presence of mixed cellularity was documented in all three ALO lines (see multiple additional examples in **Figure 2- figure supplement 2I**).

Finally, using qRT-PCR of various cell-type markers as a measure, we confirmed that the ALO models overall recapitulated the cell type composition in the adult lung tissues from which they were derived (**Figure 2- figure supplement 3**) and retained such composition in later passages without significant notable changes in any particular cell type (**Figure 2- figure supplement 4**). The mixed proximal and distal cellular composition of the ALO models and their degree of stability during in vitro culture was also confirmed by flow cytometry (**Figure 2- figure supplement 5**).

### Organoid cellularity resembles tissue sources in 3D cultures but differentiates in 2D cultures

To model respiratory infections such as COVID-19, it is necessary for pathogens to be able to access the apical surface. It is possible to microinject into the lumens of 3D organoids, as done previously with pathogens in the case of gut organoids^45–48^, or FITC-dextran in the case of lung organoids^49^, or carry out infection in apical-out 3D lung organoids with basal cells ^12^. However, the majority of the researchers have gained apical access by dissociating 3D organoids into single cells and plating them as 2D-monolayers ^10, 11, 31, 33, 50–52^. As in any epithelium, the differentiation of airway epithelial cells relies upon dimensionality (apicobasal polarity); because the loss of dimensionality can have a major impact on cellular proportions and impact disease-modeling in unpredictable ways, we assessed the impact of the 3D-to-2D conversion on cellularity by RNA seq analyses. Two commonly encountered methods of growth in 2D-monolayers were tested: (i) monolayers polarized on trans-well inserts but submerged in growth media (**Fig. 3A;** (**Figure 3- figure supplement 1A-D**), and ii) monolayers were grown at the air-liquid interface (popularly known as the ‘ALI model’ ^53, 54^ for 21-days to differentiate into the mucociliary epithelium (**Fig. 3A;** (**Figure 3- figure supplement 1E-G**). The submerged 2D-monolayers had several regions of organized vacuolated-appearing spots (**Figure 3- figure supplement 1D**; arrow), presumably due to morphogenesis and cellular organization even in 2D. The epithelial barrier was leakier, as determined by relatively lower trans-epithelial electrical resistance (TEER; **Figure 3- figure supplement 1B**) and the flux of FITC-dextran from apical to basolateral chambers (**Figure 3- figure supplement 1C**), and corroborated by morphological assessment by confocal immunofluorescence of localization of occludin, a bona-fide TJ marker. We chose occludin because it is a shared and constant marker throughout the airway that stabilizes claudins and regulates their turnover^55^ and plays an important role in maintaining the integrity of the lung epithelial barrier^56^. Junction-localized occludin was patchy in the monolayer, despite the fact that the monolayer was otherwise intact, as determined by phalloidin staining (**Figure 3- figure supplement 1H-I**). Our finding that ALO 3D organoids differentiating into monolayers in submerged cultures (where alveolar differentiation and cell-flattening happens dynamically as progenitor cells give rise to AT1/2 cells) are leaky is in keeping with prior work demonstrating that the TJs are rapidly remodeled as alveolar cells mature^57, 58^. By contrast, and as expected^59^, the ALI-monolayers formed a more effective epithelial barrier, as determined by TEER (**Figure 3- figure supplement 1F**) and appeared to be progressively hazier with time after air-lift, likely due to the accumulation of secreted mucin (**Figure 3- figure supplement 1G**).

**Figure 3.**
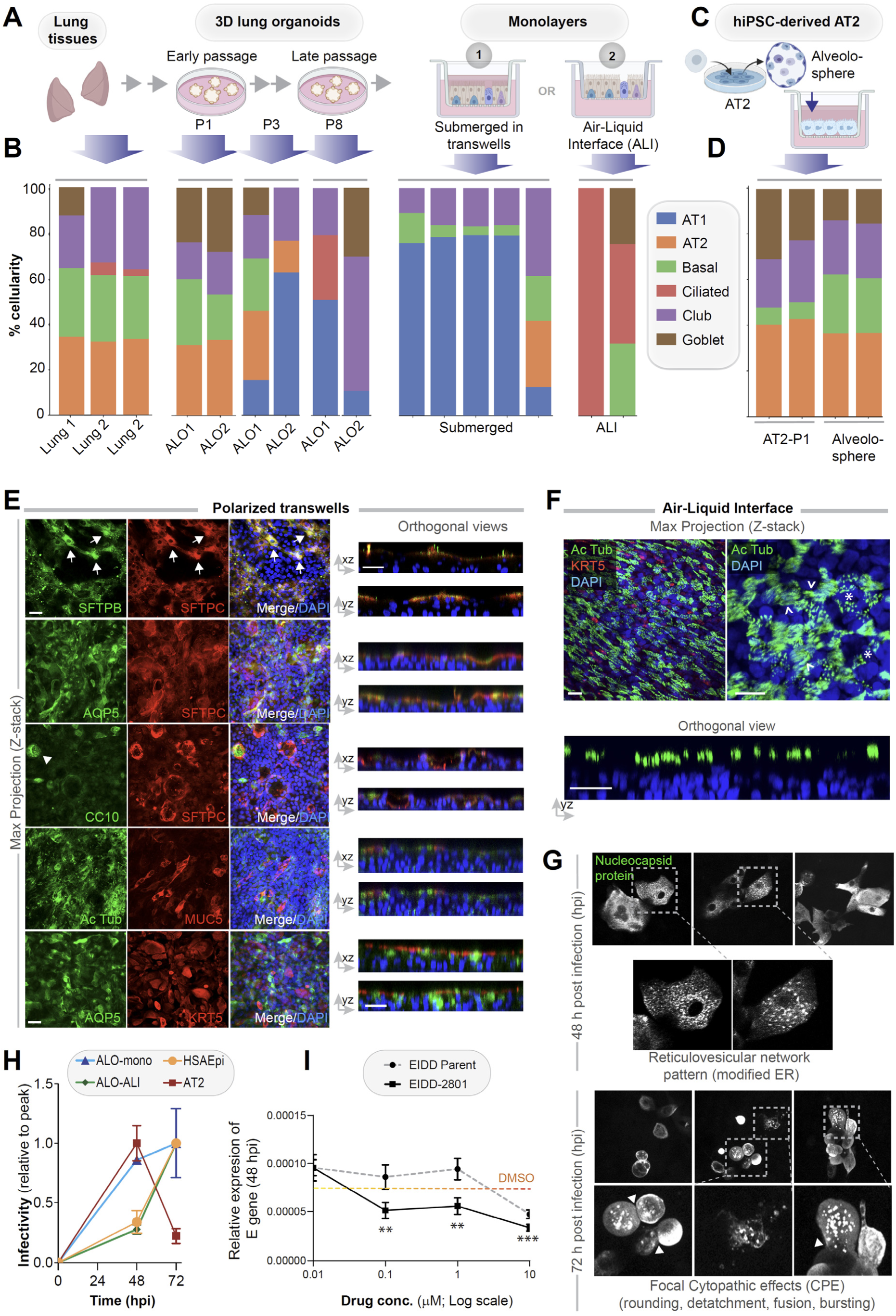
**Monolayers derived from lung organoids differentiate into proximal and distal airway components**. **A-B**. Samples collected at various steps of lung organoid isolation and expansion in culture, and from the two types of monolayers prepared using the lung organoids were analyzed by RNA seq and the datasets were compared for % cellular composition. Schematic in A shows the steps, and bar plots in B show the relative proportion of various lung cell types. GLL = general lung lineage; VEM = viral entry markers (ACE2 and TMPRSS2). **C-D**. hiPSC-derived AT2 cells and alveolospheres (C) were plated as monolayers and analyzed by RNA seq. Bar plots in D show % cellular composition. **E-F**. Submerged ALO monolayers in transwells (E) or monolayers were grown as ALI models (F) were fixed and stained for the indicated markers and visualized by confocal immunofluorescence microscopy. The representative max projected Z-stack images (left) and the corresponding orthogonal images (right) are displayed. Arrows in E indicate AT2 cells; Arrowheads in E = Club cells; asterisk in F indicate bundles of cilia standing perpendicular to the plane of the ALI monolayers; arrowheads in F indicate bundles of cilia running parallel to the plane of the ALI monolayers. Scale bar = 20 µm. **G**. Monolayers of ALO1-3 were challenged with SARS-CoV-2 for indicated time points prior to fixation and staining for KRT5, SARS-COV2 viral nucleocapsid protein and DAPI and visualized by confocal microscopy. A montage of representative images are shown, displaying reticulovesicular network patterns and various cytopathic effects. Scale bar = 15 µm. **H.** Monolayers of ALO, hiPSC-derived AT2 cells, and other alternative models (see **Figure 3- figure supplement 1-2**) were infected or not with SARS-CoV-2 and analyzed for infectivity by qPCR (targeted amplification of viral envelope, E gene). See also **Figure 3- figure supplement 3B-C** for comparison of the degree of peak viral amplification across various models.. **I**. ALO monolayers pre-treated for 4 h with either vehicle (DMSO) control or EIDD-parent (NHC) or its metabolite EIDD-2801/MK-4482 were infected with SARS-CoV-2 and assessed at 48 hpi for infectivity as in H. Line graphs display the relative expression of E gene. Error bars display S.E.M. *p* value ** < 0.01; ***, < 0.001.

RNA Seq datasets were analyzed using the same set of cell markers, as we used in **Fig. 1A** (listed in **Table 2**). Consistent with our morphologic, gene expression and FACS-based studies showcased earlier (**Fig 2; Figure 2- figure supplement 2-5**), cell-type deconvolution of our transcriptomic dataset using CIBERSORTx (https://cibersortx.stanford.edu/runcibersortx.php) confirmed that cellular composition in the human lung tissues was reflected in the 3D ALO models and that such composition was also relatively well-preserved over several passages (**Fig. 3B**; *left*); both showed a mixed population of simulated alveolar, basal, club, ciliated and goblet cells. When 3D organoids were dissociated and plated as 2D monolayers on transwells, the AT2 signatures were virtually abolished with a concomitant and prominent emergence of AT1 signatures, suggesting that growth in 2D-monolayers favor differentiation of AT2 cells into AT1 cells ^60^(**Fig. 3B**; *middle*). A compensatory reduction in proportion was also observed for the club, goblet and ciliated cells. The same organoids, when grown in long-term 2D culture conditions in the ALI model, showed a strikingly opposite pattern; alveolar signatures were almost entirely replaced by a concomitant increase in ciliated and goblet cells (**Fig. 3B**; *right*). These findings are consistent with the well-established notion that ALI conditions favor growth as pseudo-stratified mucociliary epithelium^53, 54^. As an alternative model for use as monolayers for viral infection, we developed hiPSC-derived AT2 cells and alveolospheres (**Fig. 3C**), using established protocols ^51^. Because they were grown in the presence of CHIR99021 (an aminopyrimidine derivate that is a selective and potent Wnt agonist) ^27, 28, 61^, which probably inhibits the AT2→AT1 differentiation, these monolayers were enriched for AT2 and devoid of AT1 cells (**Fig. 3D**).

The multicellularity of lung organoid monolayers was also confirmed by immunofluorescence staining and confocal microscopy of the submerged and ALI monolayers, followed by the visualization of cell markers in either max-projected z-stacks (**Fig. 3E**; *left*) or orthogonal views of the same (**Fig. 3E**; *right*). As expected, markers for the same cell type (i.e., SFTPB and SFTPC, both AT2 markers) colocalize, but markers for different cell types do not. Submerged monolayers showed the prominent presence of both AT1 (AQP5-positive) and AT2 cells. Compared to the submerged monolayers, the ALI model showed a significant increase in the ciliated epithelium (as determined by Ac Tub; compare Ac Tub stained panels in **Fig. 3E** with **3F**). This increase was associated with a concomitant decrease in KRT5-stained basal cells (**Fig. 3F**). Such loss of the basal cell marker KRT5 between submerged monolayers and the ALI model can be attributed to and the expected conversion of basal cells to other cell types (i.e., ciliated cells)^62, 63^. The presence of AT2 cells, scattered amidst the ciliated cells in these ALI monolayers, was confirmed by co-staining them for SFTPC and Ac-Tub (**Fig 3- figure supplement 1J**).

Finally, we sought to confirm that the epithelial barrier that is formed by the submerged monolayers derived from ALO are responsive to infections. To this end, we simulated infection by challenging ALO monolayers with LPS. Compared to unchallenged controls, the integrity of the barrier was impaired by LPS, as indicated by a significant drop in the TEER (**Figure 3- figure supplement 1K-L**), which is in keeping with the known disruptive role of LPS on the respiratory epithelium^64^.

Taken together, the immunofluorescence images are in agreement with the RNA seq dataset; both demonstrate that the short-term submerged monolayer favors distal differentiation (AT2→AT1), whereas the 21- day ALI model favors proximal mucociliary differentiation. It is noteworthy that these distinct differentiation phenotypes originated from the same 3D-organoids despite the seeding of cells in the same basic media composition (i.e., PneumaCult^TM^) prior to switching over to an ALI-maintenance media for the prolonged growth at Air-Liquid interface; the latter is a well-described methodology that promotes differentiation into ciliated and goblet cells ^59^.

### Differentiated 2D-monolayers show that SARS-CoV-2 infectivity is higher in proximal than distal epithelia

Because the lung organoids with complete proximodistal cellularity could be differentiated into either proximal-predominant monolayers in submerged short-term cultures or distal-predominant monolayers in long-term ALI cultures, this provided us with an opportunity to model the respiratory tract and assess the impact of the virus along the entire proximal-to-distal gradient. We first asked if ALO monolayers are permissive to SARS-CoV-2 infection and replication. Confocal imaging of infected ALO monolayers with anti-SARS-COV-2 nucleocapsid protein antibody showed that submerged ALO monolayers did indeed show progressive changes during the 48 to 72h window after infection (**Fig. 3G**): by 48 hpi we observed the formation of ‘reticulovesicular patterns’ that are indicative of viral replication within modified host endoplasmic reticulum ^65^ (**Fig. 3G**; *left*), and by 72 hpi we observed focal cytopathic effect (CPE) ^66^ such as cell-rounding, detachment, and bursting of virions (**Fig. 3G**; *right*, **Figure 3- figure supplement 3A**).

We next asked how viral infectivity varies in the various lung models. Because multiple groups have shown the importance of the ciliated airway cells for infectivity (i.e., viral entry, replication and apical release^37, 67–69^), as positive controls, we infected monolayers of human airway epithelia (see legend, **Figure 3- figure supplement 2A-D**). AT2 cells, which express high levels of viral entry receptors ACE2 and TMPRSS2 (**Fig. 1A**, **Figure 1- figure supplement 1A**) have been shown to be proficient in viral entry, but are least amenable to sustained viral release and infectivity^37, 69^. To this end, we infected monolayers of hiPSC-derived homogeneous cultures of AT2 cells as secondary controls (see legend, **Figure 3- figure supplement 2E-G**). Infection was carried out using the Washington strain of SARS-CoV-2, USA-WA1/2020 [BEI Resources NR-52281 ^70^]. As expected, the 2D-lung monolayers we generated, both the submerged or the ALI models, were readily infected with SARS-CoV-2 (**Figure 3- figure supplement 3B**), as determined by the presence of the viral envelope gene (E-gene; **Fig. 3H**); however, the kinetics of viral amplification differed. When expressed as levels of E gene normalized to the peak values in each model (**Fig. 3H**), the kinetics of the ALI-monolayer model mirrored that of the primary airway epithelial monolayers; both showed slow beginning (0 – 48 hpi) followed by an exponential increase in E gene levels from 48 to 72 hpi. The submerged monolayer model showed sustained viral infection during the 48-72 hpi window (**Figure 3- figure supplement 3B;** *left*). In the case of AT2 cells, the 48-72 hpi window was notably missing in monolayers of hiPSC-derived AT2 cells (**Fig. 3H**; **Figure 3- figure supplement 3B;** *right*). When we specifically analyzed the kinetics of viral E gene expression during the late phase (48-72 hpi window), we found that proximal airway models [human Bronchial airway Epi (HBEpC)] were more permissive than distal models [human Small Airway Epi (HSAEpC) and AT2] to viral replication (**Figure 3- figure supplement 3C**); the ALO monolayers showed intermediate sustained infectivity (albeit with variability). All models showed extensive cell death and detachment by 96 hrs and, hence, were not analyzed. Finally, using the E gene as a readout, we asked if ALO models could be used as platforms for pre-clinical drug screens. As a proof-of-concept, we tested the efficacy of nucleoside analog N^4^-hydroxycytidine (NHC; EIDD-parent) and its derivative pro-drug, EIDD-2801; both have been shown to inhibit viral replication, *in vitro* and in SARS-CoV-2- challenged ferrets^71, 72^. ALO monolayers plated in 384-wells were pre-treated for 4 h with the compounds or DMSO (control) prior to infection and assessed at 48 hpi for the abundance of E gene in the monolayers. Both compounds effectively reduced the viral titer in a dose-dependent manner (**Fig. 3I**), and the pro-drug derivative showed a better efficacy, as shown previously.

Taken together, these findings show that sustained viral infectivity is best simulated in monolayers that resemble the proximal mucociliary epithelium, i.e., 2D-monolayers of lung organoids grown as ALI models and the primary airway epithelia. Because prior studies conducted in patient-derived airway cells ^37^ mirrors what we see in our monolayers, we conclude that proximal airway cells within our mixed-cellular model appear to be sufficient to model viral infectivity in COVID-19. Findings also provide proof-of-concept that ALO monolayers may be adapted in miniaturized formats for use in 384-well plates for high-throughput (HTP) drug screens.

### Differentiated 2D-monolayers show that host immune response is higher in distal than proximal epithelia

Next, we asked if the newly generated lung models accurately recapitulate the host immune response in COVID-19. To this end, we analyzed the infected ALO monolayers (both the submerged and ALI variants) as well as the airway epithelial (HSAEpC) and AT2 monolayers by RNA seq and compared them all against the transcriptome profile of lungs from deceased COVID-19 patients. We did this analysis in two steps of reciprocal comparisons: (i) First, the actual human disease-derived gene signature was assessed for its ability to distinguish infected from uninfected disease models (in **Fig 4**). (ii) Second, the ALO model-derived gene signature was assessed for its ability to distinguish healthy from diseased patient samples (in **Fig 5**). A publicly available dataset (GSE151764) ^73^, comprised of lung transcriptomes from victims deceased either due to non-infectious causes (controls) or due to COVID-19, was first analyzed for differentially expressed genes (**Fig. 4A-B**). This cohort was chosen as a test cohort over others because it was the largest one available at the time of this study with appropriate postmortem control samples. Differentially expressed genes showed an immunophenotype that was consistent with what is expected in viral infections (**Fig. 4C; Table 4**; **Figure 4- figure supplement 1**), and showed overrepresentation of pathways such as interferon, immune, and cytokine signaling (**Fig. 4D; Table 5**; **Figure 4- figure supplement 2**). Differentially expressed gene signatures and reactome pathways that were enriched in the test cohort were fairly representative of the host immune response observed in patient-derived respiratory samples in multiple other validation cohorts; the signature derived from the test cohort could consistently classify control (normal) samples from COVID-19-samples (ROC AUC 0.89 to 1.00 across the board; **Fig. 4E**). The most notable finding is that the patient-derived signature was able to perfectly classify the EpCAM-sorted epithelial fractions from the bronchoalveolar lavage fluids of infected and healthy subjects (ROC AUC 1.00; GSE145926-Epithelium ^74^, suggesting that the respiratory epithelium is a major site where the host immune response is detected in COVID-19. When compared to existing organoid models of COVID-19, we found that the patient-derived COVID-19-lung signature was able to perfectly classify infected *vs*. uninfected late passages (> 50) of hiPSC-derived AT1/2 monolayers (GSE155241) ^50^ and infected *vs.* uninfected liver and pancreatic organoids (**Fig. 4F**). The COVID-19-lung signatures failed to classify commonly used respiratory models, e.g., A549 cells and bronchial organoids, as well as intestinal organoids (**Fig. 4F**). A similar analysis on our own lung models revealed that the COVID-19-lung signature was induced in submerged monolayers with distal-predominant AT2→AT1 differentiation but not in the proximal-predominant ALI model (ROC AUC 1.00 and 0.50, respectively; **Fig. 4G**). The ALI model and the small airway epithelia, both models that mimic the airway epithelia (and lack alveolar pneumocytes; see **Fig. 3B**), failed to mount the patient-derived immune signatures (**Fig. 4H**; *left*). These findings suggested that the presence of alveolar pneumocytes is critical for emulating host response. To our surprise, induction of the COVID-19-lung signature also failed in hiPSC-derived AT2 monolayers (**Fig.4H**; *right*), indicating that AT2 cells are unlikely to be the source of such host response. These findings indicate that both proximal airway and AT2 cells, when alone, are insufficient to induce the host immune response that is encountered in the lungs of COVID-19 patient.

**Figure 4.**
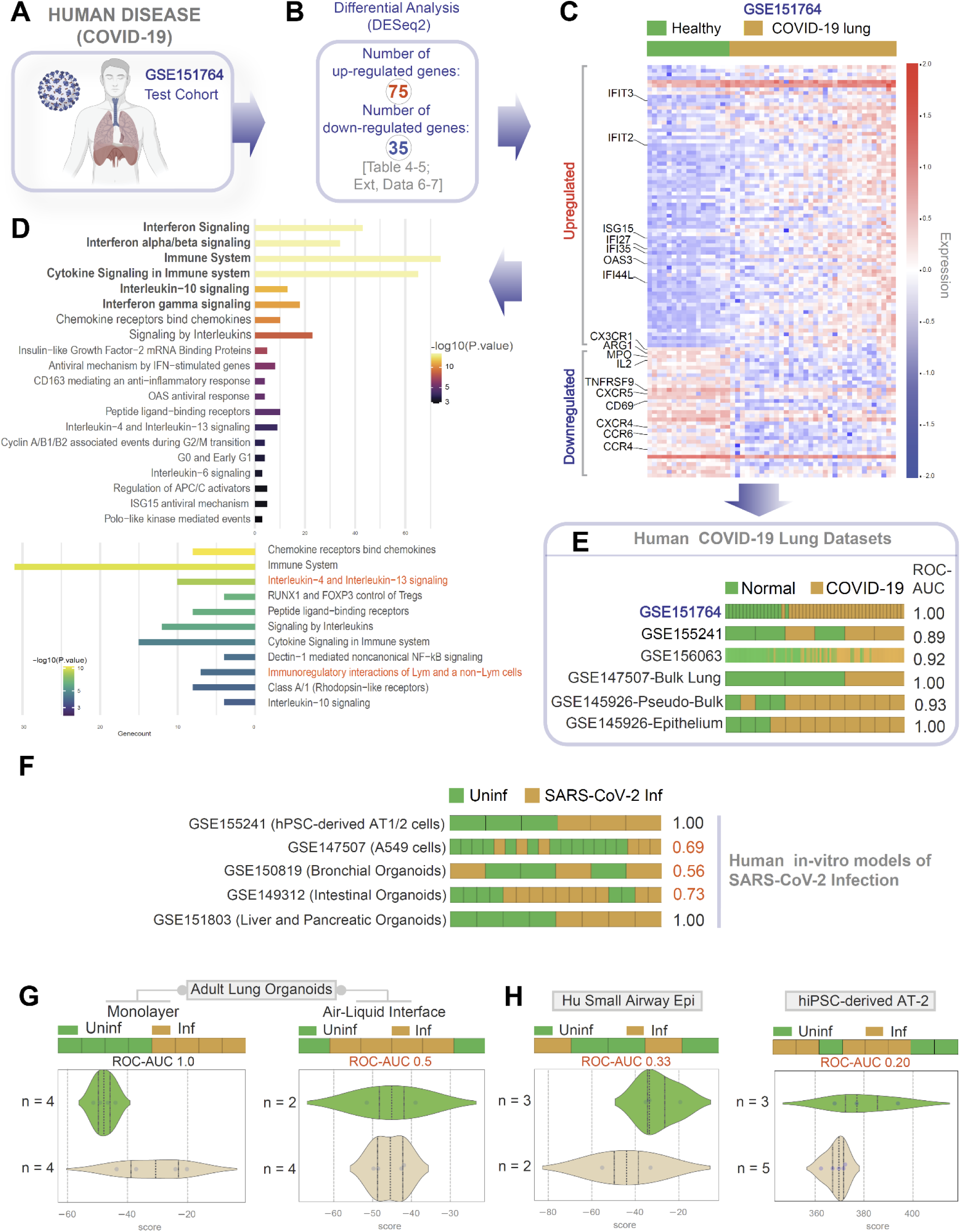
**Gene expression patterns in the lungs of patients with COVID-19 (actual disease) is recapitulated in lung organoid monolayers infected with SARS-CoV-2 (disease model)**. **A-C**. Publicly available RNA Seq datasets (GSE151764) of lung autopsies from patients who were deceased due to COVID-19 or non-infectious causes (healthy normal control) were analyzed for differential expression of genes (B). The DEGs are displayed as a heatmap labeled with selected genes in C. See also **Figure 4- figure supplement 1** for the same heatmap with all genes labeled. **D**. Reactome-pathway analysis shows the major pathways up- or downregulated in the COVID-19-afflicted lungs. See also **Figure 4- figure supplement 2** for visualization as hierarchical ReacFoam. **E**. Bar plots display the ability of the differentially expressed genes in the test cohort (GSE151764) to classify human COVID-19 respiratory samples from four other independent cohorts. **F**. Bar plots display the ability of the differentially expressed genes in the test cohort (GSE151764) to classify published *in vitro* models for SARS-CoV-2 infection where RNA Seq datasets were either generated in this work or publicly available. **G**-**H.** Bar (top) and violin (bottom) plots compare the relative accuracy of disease modeling in four *in vitro* models used in the current work, as determined by the induction of COVID-19 lung signatures in each model. G; monolayer (left) and ALI models (right) prepared using adult lung organoids (ALOs). H; Primary human small airway epithelium (left) and hiPSC –derived AT2-monolayers (right). **Table 8** lists details regarding the patient cohorts/tissue or cell types represented in each transcriptomic dataset.

**Figure 5.**
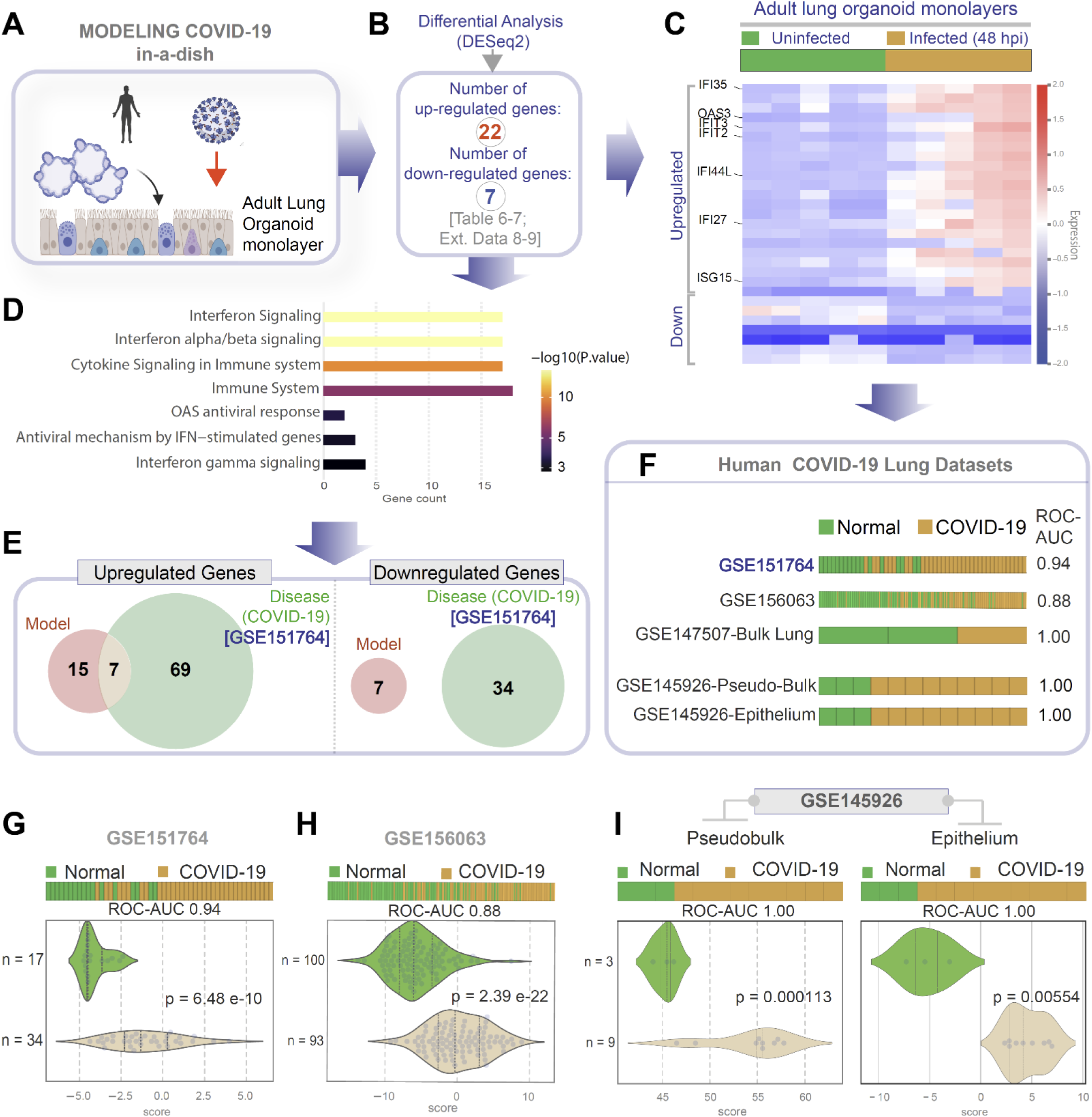
**Genes and pathways induced in the SARS-CoV-2-infected lung organoid monolayers (disease model) are induced also in the lungs of COVID-19 patients (actual disease). A-C**. Adult lung organoid monolayers infected or not with SARS-CoV-2 were analyzed by RNA seq and differential expression analysis. Differentially expressed genes (B) are displayed as a heatmap in C. While only selected genes are labeled in panel C (which represent overlapping DEGS between our organoid model and publicly available COVID-19 lung dataset, GSE151764), the same heatmap is presented in **Figure 5- figure supplement 1** with all genes labeled. **D.** Reactome-pathway analysis shows the major pathways upregulated in SARS-CoV-2-infected lung organoid monolayers. See also **Figure 5- figure supplement 2** for visualization as hierarchical ReacFoam. **E**. A Venn diagram showing overlaps in DEGs between model (current work; 5B) and disease (COVID-19 lung dataset, GSE151764; Fig 4). **F**. Bar plots display the ability of the DEGs in infected lung monolayers to classify human normal vs COVID-19 respiratory samples from five independent cohorts. **G-I**. Bar (top) and violin (bottom) plots compare the accuracy of disease modeling in three publicly available human lung datasets, as determined by the significant induction of the DEGs that were identified in the SARS-CoV-2-challenged monolayers. See also **Table 8**, which enlists details regarding the patient cohorts/tissue or cell types represented in each transcriptomic dataset.

**Table 4:** Upregulated genes and pathways: Healthy vs COVID-19 lung (<u>GSE151764</u>)

|  |  |  |  |  |
| --- | --- | --- | --- | --- |
| BRCA2<br>CYBB<br>KRT5<br>C1QB<br>FCGR1A<br>IL10<br>IL6<br>CD44<br>CD276 | XAGE1B<br>CCR5<br>CCR2<br>ALOX15B<br>CMKLR1<br>MX1<br>TNFRSF17<br>CCR1<br>CXCR3 | CDK1<br>GBP1<br>HLA-G<br>IDO1<br>ISG20<br>LAG3<br>MAD2L1<br>CXCL9<br>MKI67 | SNAI2<br>IFITM1<br>GZMB<br>CD163<br>CD38<br>BST2<br>BUB1<br>CCL20<br>CCNB2 | CXCL11<br>IFI27<br>IFI35<br>TDO2<br>GZMA<br>OAS3<br>POU2AF1<br>CXCL13<br>GNLY |
| DMBT1<br>DDX58<br>TNFAIP8<br>LAMP3<br>KIAA0101<br>MELK | SLAMF8<br>IL21<br>FOXM1<br>IFIH1<br>IFI6<br>PDCD1LG2 | IFIT2<br>IFIT1<br>CXCL10<br>IRF4<br>PSMB9<br>CCL18 | TNFSF18<br>ISG15<br>CDKN3<br>C1QA<br>OAS1<br>OAS2 | IFIT3<br>TOP2A<br>LILRB1<br>HERC6<br>TNFSF13B<br>IFI44L<br>STAT1 |

**Pathways:**
| Name | pValue | FDR |
| --- | --- | --- |
| Interferon Signaling | 1.1102230246251565e-16 | 1.1102230246251565e-14 |
| Interferon alpha/beta signaling | 1.1102230246251565e-16 | 1.1102230246251565e-14 |
| Cytokine Signaling in Immune system | 1.1102230246251565e-16 | 1.1102230246251565e-14 |
| Immune System | 1.1102230246251565e-16 | 1.1102230246251565e-14 |
| Interleukin-10 signaling | 9.845457782375888e-13 | 7.87636622590071e-11 |
| Interferon gamma signaling | 9.256706512417168e-12 | 6.109426298195331e-10 |
| Chemokine receptors bind chemokines | 1.0827139185209944e-10 | 6.171469335569668e-9 |
| Signaling by Interleukins | 6.813459352272844e-9 | 3.406729676136422e-7 |
| Insulin-like Growth Factor-2 mRNA Binding Proteins (IGF2BPs/IMPs/VICKZs) bind RNA | 1.268436958801189e-7 | 0.000005581122618725232 |
| Antiviral mechanism by IFN-stimulated genes | 0.000001933058349390393 | 0.00007732233397561572 |
| CD163 mediating an anti-inflammatory response | 0.000007798676168513374 | 0.0002807523420664815 |
| OAS antiviral response | 0.000010208709973369423 | 0.00033688742912119096 |
| Peptide ligand-binding receptors | 0.000017140576871854662 | 0.0005142173061556399 |
| Interleukin-4 and Interleukin-13 signaling | 0.00010149486613275638 | 0.0028418562517171786 |
| Cyclin A/B1/B2 associated events during G2/M transition | 0.00018878164653901575 | 0.004908322810014409 |
| G0 and Early G1 | 0.00036071218383249004 | 0.009017804595812251 |
| Interleukin-6 signaling | 0.0004656678443889506 | 0.010710360420945864 |
| ISG15 antiviral mechanism | 0.0008313991987842773 | 0.017459383174469822 |
| Regulation of APC/C activators between G1/S and early anaphase | 0.0008313991987842773 | 0.017459383174469822 |
| Polo-like kinase mediated events | 0.0011105065129394243 | 0.022210130258788485 |
| APC/C-mediated degradation of cell cycle proteins | 0.0013081035810655894 | 0.02354586445918061 |
| Regulation of mitotic cell cycle | 0.0013081035810655894 | 0.02354586445918061 |
| G2/M DNA replication checkpoint | 0.001750156331829622 | 0.029752657641103575 |
| Class A/1 (Rhodopsin-like receptors) | 0.0023550630454979293 | 0.035376667816378604 |
| Interleukin-6 family signaling | 0.002358444521091907 | 0.035376667816378604 |
| TNFs bind their physiological receptors | 0.002358444521091907 | 0.035376667816378604 |

**Table 5:** Downregulated genes and pathways: Healthy vs COVID-19 lung (<u>GSE151764</u>)

|  |  |  |  |  |
| --- | --- | --- | --- | --- |
| CX3CR1<br>ARG1<br>MPO<br>IL2<br>BCL2<br>CA4<br>IGF1R | JAML<br>CX3CR1<br>HLA-DQB2<br>TNFRSF9<br>CXCR5<br>CD1C<br>CD69 | KLRB1<br>LY9<br>CCL17<br>CCL22<br>TCF7<br>CXCR4<br>CD83 | GRAP2<br>MMP9<br>RORC<br>CCR4<br>IRS1<br>ITK<br>KLRG1 | CD226<br>CD160<br>FOXP3<br>CRTAM<br>CCR6<br>CEACAM8<br>PTGS2 |

**Pathways:**
| Name | pValue | FDR |
| --- | --- | --- |
| Chemokine receptors bind chemokines | 2.8463564838432376e-11 | 4.981123846725666e-9 |
| Immune System | 1.2524759007703778e-10 | 1.0896540336702287e-8 |
| Interleukin-4 and Interleukin-13 signaling | 2.820398359304477e-9 | 1.6358310483965965e-7 |
| RUNX1 and FOXP3 control the development of regulatory T lymphocytes (Tregs) | 4.3109720904155324e-7 | 0.00001853717998878679 |
| Peptide ligand-binding receptors | 6.709444980712576e-7 | 0.000023483057432494014 |
| Signaling by Interleukins | 0.0000015036584932737185 | 0.000043606096304937836 |
| Cytokine Signaling in Immune system | 0.000026065058545632347 | 0.0006516264636408087 |
| Dectin-1 mediated noncanonical NF-kB signaling | 0.00008640543214522012 | 0.0018145140750496225 |
| Immunoregulatory interactions between a Lymphoid and a non-Lymphoid cell | 0.00010833886747929622 | 0.002058438482106628 |
| Class A/1 (Rhodopsin-like receptors) | 0.0001833048828232542 | 0.0031161830079953212 |
| Interleukin-10 signaling | 0.0002366961933546463 | 0.0035504429003196947 |
| RUNX3 Regulates Immune Response and Cell Migration | 0.0005791814112650062 | 0.0077471849338778265 |
| Extra-nuclear estrogen signaling | 0.0005959373026059867 | 0.0077471849338778265 |
| BH3-only proteins associate with and inactivate anti-apoptotic BCL-2 members | 0.000699254752329681 | 0.008391057027956172 |
| CLEC7A (Dectin-1) signaling | 0.000822803514531345 | 0.009050838659844795 |
| Generation of second messenger molecules | 0.0011719919081142338 | 0.011719919081142338 |
| Innate Immune System | 0.0016764040917021106 | 0.015723603669013264 |
| GPCR ligand binding | 0.0017470670743348071 | 0.015723603669013264 |
| Adaptive Immune System | 0.002059835990545711 | 0.018538523914911398 |
| Estrogen-dependent nuclear events downstream of ESR-membrane signaling | 0.004670055830237274 | 0.03736044664189819 |
| C-type lectin receptors (CLRs) | 0.005458044949850915 | 0.04366435959880732 |
| Transcriptional regulation by RUNX3 | 0.008124332598870021 | 0.05687032819209015 |
| BMAL1:CLOCK,NPAS2 activates circadian gene expression | 0.009518272708583853 | 0.06662790896008697 |
| ESR-mediated signaling | 0.012073762373888686 | 0.0845163366172208 |
| Transcriptional regulation by RUNX1 | 0.012881563708187715 | 0.08786708746501692 |
| TCR signaling | 0.01464451457750282 | 0.08786708746501692 |

**Table 8.** The list of GSE numbers used in the figures.

| **GSE #** | **Cell Type/ Tissue** | **References** | **Figure** |
| --- | --- | --- | --- |
| **GSE132914** | Tissue from Idiopathic pulmonary fibrosis subjects and donor controls | PMID: 32991815 | 1A |
| **GSE151764** | COVID-19 and normal lung tissue post-mortem | PMID: 33033248 | 4A-4E, 5E-5G |
| **GSE155241** | hPSC lung organoids and colon organoids infected with SARS-CoV-2 | PMID: 33116299 | 4E-4F, 6D |
| **GSE156063** | Upper airway of COVID-19 patients and other acute respiratory illnesses | PMID: 33203890 | 4E, 5F, 5H |
| **GSE147507** | A549 cells and bulk lung | PMID: 32416070; PMID: 33782412 | 4E- 4F, 5F |
| **GSE145926** | Bronchoalveolar lavage fluid (BALF) immune cells from COVID-19 and healthy subjects | PMID: 32398875 | 4E, 5F, 5I |
| **GSE150819** | Human bronchial organoids | From commercially available HBEpC, | 4F, 6C |
| **GSE149312** | Intestinal organoids infected with SARS-CoV or SARS-CoV-2 | PMID: 32358202 | 4F |
| **GSE151803** | hPSC derived pancreatic and lung organoids infected with SARS-CoV-2 | no publication yet | 4F |
| **GSE153940** | Vero E6 control or SARS-CoV-2 infected cells | PMID: 32707573 | 6B |
| **GSE153218** | SARS-CoV-2 infected bronchoalveolar cells derived from organoids grown using progenitor cells from human fetal lung but tip (LBT). | PMID: 33283287 | 6H |

Next, we analyzed the datasets from our ALO monolayers for differentially expressed genes (DEGs) when challenged with SARS-COV-2 (**Fig. 5A-B**). Genes and pathways upregulated in the infected lung organoid-derived monolayer models (**Figure 5- figure supplement 1-2**) overlapped significantly with those that were upregulated in the COVID-19 lung signature (compare **Fig. 4C-D** with **5C-D**, **Table 6-7**). We observed only a partial overlap (rangng from ∼22-55% across various human datasets; **Figure 5-Figure Supplement 3**) in upregulated genes and no overlaps among downregulated genes between model and disease (COVID-19) (**Fig. 5E**). Because the degree of overlap was even lesser (ranging from ∼10-25% across various human datasets; **Figure 5-Figure Supplement 3**) in the case of another publicly released model (GSE160435)^11^, these discrepancies between the model and the actual disease likely reflect the missing stromal and immune components in our organoid monolayers. Regardless of these missing components, the model-derived DEG signature was sufficient to consistently and accurately classify diverse cohorts of patient-derived respiratory samples (ROC AUC ranging from 0.88 to 1.00; **Fig. 5F**); the model-derived DEG signature was significantly induced in COVID-19 samples compared to normal controls (**Fig. 5G-H**). Most importantly, the model-derived DEG signature was significantly induced in the epithelial cells recovered from bronchoalveolar lavage (**Fig. 5I**).

**Table 6:** Upregulated genes and pathways: Uninfected vs Infected (48 hpi) lung organoid monolayers

|  |  |  |  |
| --- | --- | --- | --- |
| IFI35 | EPSTI1 | AMIGO2 | IFITM2 |
| SLC4A11 | CMPK2 | WARS1 | FAAP100 |
| APOL1 | OASL | IFI27 | ISG15 |
| OAS3 | IFI44L | CD14 | SLC35F6 |
| IFIT3 | IFI44 | SAMD9L |  |
| IFIT2 | PARP9 | SRP9P1 |  |

**Pathways:**
| Name | pValue | FDR |
| --- | --- | --- |
| Interferon Signaling | 1.1102230246251565e-16 | 4.218847493575595e-15 |
| Interferon alpha/beta signaling | 1.1102230246251565e-16 | 4.218847493575595e-15 |
| Cytokine Signaling in Immune system | 1.1540801647669241e-10 | 2.8852004119173102e-9 |
| Immune System | 0.000002540114879323596 | 0.00004826218270714833 |
| OAS antiviral response | 0.0004764545663257236 | 0.007146818494885854 |
| Antiviral mechanism by IFN-stimulated genes | 0.0010333472605504879 | 0.012400167126605854 |
| Interferon gamma signaling | 0.0018896946191467867 | 0.020786640810614654 |
| Transfer of LPS from LBP carrier to CD14 | 0.006318772245420878 | 0.056868950208787905 |
| TRIF-mediated programmed cell death | 0.020912675855586538 | 0.16560733288455465 |
| MyD88 deficiency (TLR2/4) | 0.03733748270572135 | 0.16560733288455465 |
| IRAK2 mediated activation of TAK1 complex upon TLR7/8 or 9 stimulation | 0.03733748270572135 | 0.16560733288455465 |
| TRAF6-mediated induction of TAK1 complex within TLR4 complex | 0.03937173812023953 | 0.16560733288455465 |
| IRAK4 deficiency (TLR2/4) | 0.03937173812023953 | 0.16560733288455465 |
| Activation of IRF3/IRF7 mediated by TBK1/IKK epsilon | 0.04140183322113866 | 0.16560733288455465 |
| Caspase activation via Death Receptors in the presence of ligand | 0.04140183322113866 | 0.16560733288455465 |
| IKK complex recruitment mediated by RIP1 | 0.04948077475965562 | 0.18550132647663609 |

**Table 7:** Downregulated genes and pathways: Uninfected vs Infected (48 hpi) lung organoid monolayers

|  |  |  |  |
| --- | --- | --- | --- |
| AC093392.1 | ARHGAP19 | HLA-V | RN7SL718P |
| MT-TV | AC138969.3 | AC016766.1 |  |

**Pathways:**
| Name | pValue | FDR |
| --- | --- | --- |
| rRNA processing in the mitochondrion | 0.01892731245535506 | 0.08366120772817287 |
| tRNA processing in the mitochondrion | 0.021271491049123403 | 0.08366120772817287 |
| Mitochondrial translation termination | 0.043991554455812154 | 0.08366120772817287 |
| Mitochondrial translation elongation | 0.043991554455812154 | 0.08366120772817287 |
| Mitochondrial translation initiation | 0.04490921761868061 | 0.08366120772817287 |
| Mitochondrial translation | 0.04765767844112356 | 0.08366120772817287 |

Taken together, these cross-validation studies from disease to model (**Fig. 4**) and *vice versa* (**Fig. 5**) provide an objective assessment of the match between the host response in COVID-19 lungs and our submerged ALO monolayers. Such a match was not seen in the case of the other models, e.g., the proximal airway-mimic ALI model, HSAEpC monolayer, or hiPSC-derived AT2 models. Because the submerged ALO monolayers contained both proximal airway epithelia (basal cells) and promoted AT2→AT1 differentiation, findings demonstrate that mixed cellular monolayers can mimic the host response in COVID-19. A subtractive analysis revealed that the cell type that is shared between models which showed induction of host response signatures (i.e., ALO submerged monolayers and GSE155241 ^50^; **Fig. 5F**) but is absent in models that do not show such response (hu Bronchial organoids, small airway epi, ALI-model of ALO) is AT1. We conclude that distal differentiation from AT2→AT1, a complex process that is comprised of distinct intermediates ^75^, is essential for modeling the host immune response in COVID-19. Further experimental evidence is needed to directly confirm if and which intermediate states during the differentiation of AT2 to AT1 is essential for the immune response to COVID19.

### Both proximal and distal airway epithelia are required to mount the overzealous host response in COVID-19

We next asked which model best simulated the overzealous host immune response that has been widely implicated in fatal COVID-19. To this end, we relied upon a recently described artificial intelligence (AI)-guided definition of the nature of the overzealous response in fatal COVID-19 ^76^. Using ACE2 as a seed gene, a 166-gene signature was identified and validated as an invariant immune response that was shared among all respiratory viral pandemics, including COVID-19 (**Fig. 6A**). A subset of 20 genes within the 166-gene signature was subsequently identified as a determinant of disease severity/fatality; these 20 genes represented translational arrest, senescence, and apoptosis. These two signatures, referred to as ViP (166-gene) and severe ViP (20-gene) signatures, were used as a computational framework to first vet existing SARS-CoV-2 infection models that have been commonly used for therapeutic screens (**Fig. 6B-D**). Surprisingly, we found that each model fell short in one way or another. For example, the Vero E6, which is a commonly used cultured cell model, showed a completely opposite response; instead of being induced, both the 166-gene and 20-gene ViP signatures were suppressed in infected Vero E6 monolayers (**Fig. 6B**). Similarly, neither ViP signature was induced in the case of SARS-CoV-2 challenged human bronchial organoids^16^ (**Fig. 6C**). Finally, in the case of the hiPSC-derived AT1/2 organoids, which recapitulated the COVID-19-lung derived immune signatures (in **Fig. 4F**), the 166-gene ViP signature was induced significantly (**Fig. 6D**; *top*), but the 20-gene severity signature was not (**Fig. 6D**; *bottom*). These findings show that none of the existing models capture the overzealous host immune response that has been implicated in a fatality.

**Figure 6.**
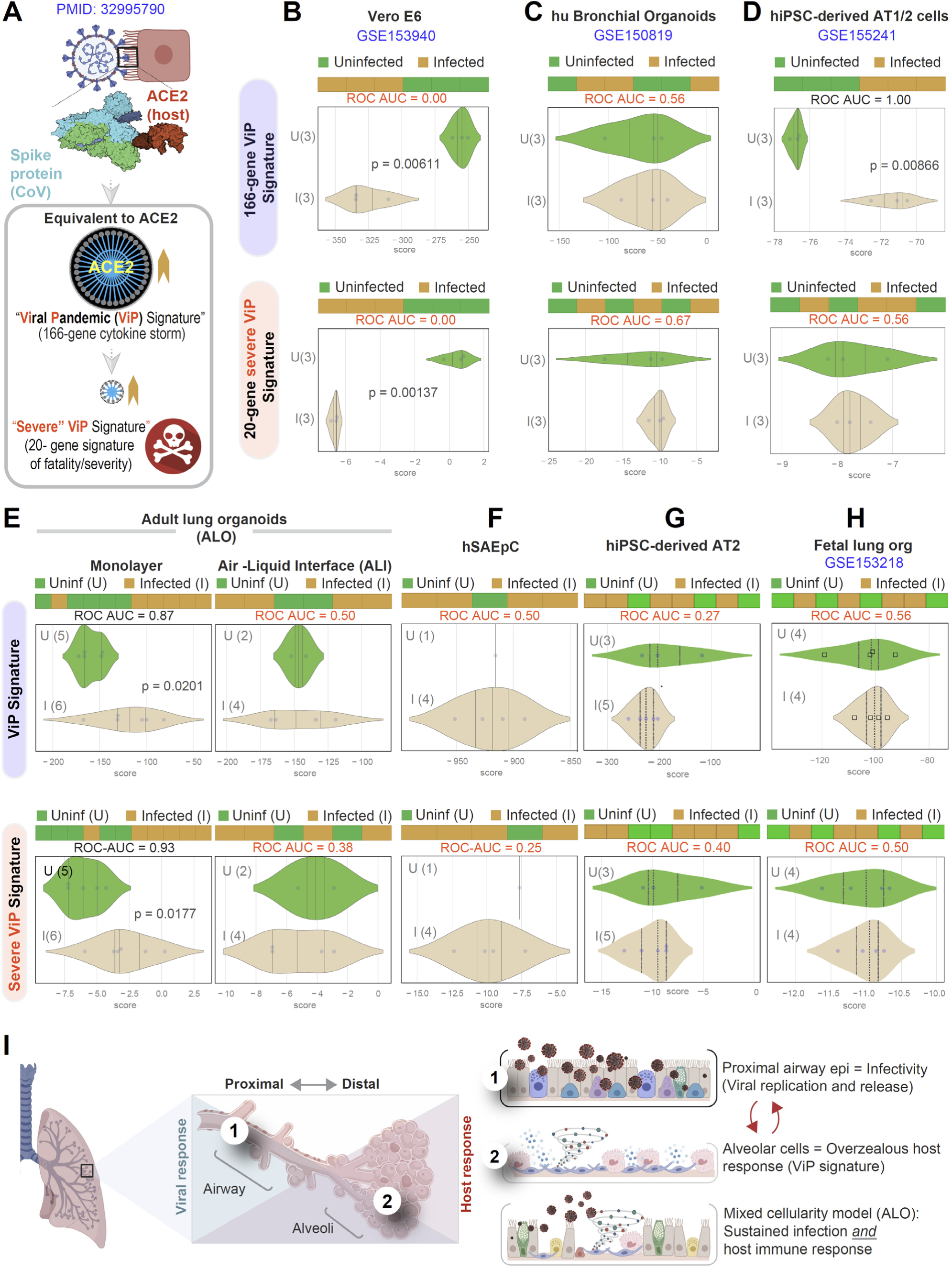
**Both proximal and distal airway components are required to model the overzealous host response in COVID-19. A.** Schematic summarizing the immune signatures identified based on ACE2-equivalent gene induction observed invariably in any respiratory viral pandemic. The 166-gene ViP signature captures the cytokine storm in COVID-19, whereas the 20-gene subset severe ViP signature is indicative of disease severity/fatality. **B-D**. Publicly available RNA Seq datasets from commonly used lung models, Vero E6 (B), human bronchial organoids (C) and hPSC-derived AT1/2 cell-predominant lung organoids are classified using the 166-gene ViP signature (top row) and 20-gene severity signature (bottom row). **E-G**. RNA Seq datasets generated in this work using either human small airway epithelial cells (E), adult lung organoids as submerged or ALI models (left and right, respectively, in F) and hiPSC-derived AT2 cells (G) were analyzed and visualized as in B-D. **H**. Publicly available RNA seq datasets from fetal lung organoid monolayers^77^ infected or not with SARS-CoV-2 were analyzed as in B-D for the ability of ViP signatures to classify infected (I) from uninfected (U) samples. ROC AUC in all figure panels indicate the performance of a classification model using the ViP signatures. **I**. Summary of findings in this work, its relationship to the observed clinical phases in COVID-19, and key aspects of modeling the same. **Table 8** lists details regarding the patient cohorts/tissue or cell types represented in each transcriptomic dataset.

Our lung models showed that both the 166- and 20-gene ViP signatures were induced significantly in the submerged ALO-derived monolayers that had distal differentiation (**Fig. 6E;** *left*), but not in the proximal-mimic ALI model (**Fig. 6E;** *right*). Neither signatures were induced in monolayers of small airway epithelial cells (**Fig. 6F**) or hiPSC-derived AT2 cells (**Fig. 6G**). Finally, we analyzed a recently published lung organoid model that that supports robust SARS-CoV-2 infection; this model was generated using multipotent SOX2+SOX9+ lung bud tip (LBT) progenitor cells that were isolated from the canalicular stage of human fetal lungs (∼16–17 wk post-conception)^77^. Despite mixed cellularity (proximal and distal), this fetal lung organoid model failed to induce the ViP signatures^76^ (**Fig 6H**). These findings indicate that despite having mixed cellular composition and the added advantage of being permissive to robust viral replication (achieving ∼ 5 log-fold increase in titers), the model lacks the signature host response that is seen in all human samples of COVID-19.

Taken together with our infectivity analyses, these findings demonstrate that although the proximal airway epithelia and AT2 cells may be infected, and as described by others ^21, 37^, may be vital for mounting a viral response and for disease transmission, these cells alone cannot mount the overzealous host immune response that is associated with the fatal disease. Similarly, even though the alveolar pneumocytes, AT1 and AT2 cells, are sufficient to mount the host immune response, in the absence of proximal airway components, they too are insufficient to recapitulate the severe ViP signature that is characterized by cellular senescence and apoptosis. However, when both proximal and distal components are present, i.e., basal, ciliated and AT1 cells, the model mimicked the overzealous host immune response in COVID-19 (**Fig. 6I**).

## Discussion

The most important discovery we report here is the creation of adult lung organoids that are complete with both proximal airway and distal alveolar epithelia; these organoids can not only be stably propagated and expanded in 3D cultures but also used as monolayers of mixed cellularity for modeling viral and host immune responses during respiratory viral pandemics. Furthermore, an objective analysis of this model and other existing SARS-CoV-2-infected lung models against patient-lung derived transcriptomes showed that the model which most closely emulates the elements of viral infectivity, lung injury, and inflammation in COVID-19 is one that contained both proximal and distal alveolar signatures (**Fig 6H**), whereas, the presence of just one or the other fell short.

There are three important impacts of this work. First, successful creation of adult human lung organoids that are complete with both proximal and distal signatures has not been accomplished before. Previous works show the successful use of airway basal cells for organoid creationtion, but ensuring the completeness of the model with all other lung cells has been challenging to create ^78^. The multicellularity of the lung has been a daunting challenge that many experts have tried to recreate *in vitro*; in fact, the demand for perfecting such a model has always remained high, not just in the current context of the COVID-19 pandemic but also with the potential of future pandemics. We have provided the evidence that the organoids that were created using our methodology retain proximal and distal cellularity throughout multiple passages and even within the same organoid. Although a systematic design of experiment (DoE) approach ^79^ was not involved in getting to this desirable goal, a rationalized approach was taken. For example, a Wnt/R-spondin/Noggin-containing conditioned media was used as a source of the so-called ‘niche factors’ for any organoid growth ^80^. This was supplemented with recombinant FGF7/10; FGF7 is known to help cell proliferation and differentiation and is required for normal branching morphogenesis ^81^, whereas FGF10 helps in cell maturation ^82^ and in alveolar regeneration upon injury^83^. Together, they are likely to have directed the differentiation toward distal lung lineages (hence, the preservation of alveolar signatures). The presence of both distal alveolar and proximal ciliated cells was critical: proximal cells were required to recreate sustained viral infectivity, and the distal alveolar pneumocytes, in particular, the ability of AT2 cells to differentiate into AT1 pneumocytes was essential to recreate the host response. It is possible that the response is mediated by a distinct AT2-lineage population, i.e., damage-associated transient progenitors (DATPs), which arise as intermediates during AT2→AT1 differentiation upon injury-induced alveolar regeneration ^75^. Although somewhat unexpected, the role of AT1 pneumocytes in mounting innate immune responses has been documented before in the context of bacterial pneumonia ^84, 85^. In work ^51^ that was published during the preparation of this manuscript, authors used long-term ALI models of hiPSC-derived AT2 monolayers (in growth conditions that inhibit AT2→AT1 differentiation, as we did here for our AT2 model) and showed that SARS-CoV-2 induces iAT2-intrinsic cytotoxicity and inflammatory response, but failed to induce type 1 interferon pathways (IFN α/β). It is possible that prolonged culture of iAT2 pneumocytes gives rise to some DATPs but cannot robustly do so in the presence of inhibitors of AT1 differentiation. This (spatially segregated viral and host immune response) is a common theme across many lung infections (including bacterial pneumonia and other viral pandemics ^37, 86–88^ and hence, this mixed cellularity model is appropriate for use in modeling diverse lung infections and respiratory pandemics to come.

Second, among all the established lung models so far, ours features 4 key properties that are desirable whenever disease models are being considered for their use in HTP modes for rapid screening of candidate therapeutics and vaccines —(i) reproducibility, propagability and scalability, (ii) cost-effectiveness, (iii) personalization, and (iv) modularity, with the potential to add other immune and non-immune cell types to our multi-cellular complex lung model. We showed that the protocol we have optimized supports isolation, expansion and propagability at least up to 12-15 passages (at the time of submission of this work), with documented retention of proximal and distal airway components up to P8 (by RNA seq). Feasibility has also been established for scaling up and optimizing the conditions for them to be used in miniaturized 384-well infectivity assays. We also showed that the protocols for generating lung organoids could be reproduced in both genders and regardless of the donor’s smoking status, consistency in outcome and growth characteristics were observed across all isolation attempts. The ALOs are also cost-effective; the need for exclusive reliance on recombinant growth factors was replaced at least in part with conditioned media from a commonly used cell line (L-WRN/ ATCC® CRL-2647 cells). Such media has a batch to batch stable cocktail of Wnt, R-Spondin, and Noggin and has been shown to improve reproducibility in the context of GI organoids in independent laboratories ^89^. By that token, our culture conditions may have also improved reproducibility. The major disadvantage, however, remains that the composition of the media is undefined. Because the model is propagable, repeated iPSC-reprogramming (another expensive step) is also eliminated, further cutting costs compared to many other models. As for personalization, our model is derived from adult lung stem cells from deep lung biopsies; each organoid line was established from one patient. By avoiding iPSCs or EPSCs, this model not only captures genetics but also retains organ-specific epigenetic programming in the lung, and hence, potentially additional programming that may occur in disease (such as in the setting of chronic infection, injury, inflammation, somatic mutations, etc.). The ability to replicate donor phenotype and genotype in vitro allows for potential use as pre-clinical human models for Phase ‘0’ clinical trials. As for modularity, by showing that the 3D lung organoids could be used as polarized monolayers on transwells to allow infectious agents to access the apical surface (in this case, SARS-CoV-2), we demonstrate that the organoids have the potential to be reverse-engineered with additional components in a physiologically relevant spatially segregated manner: for example, immune and stromal cells can be placed in the lower chamber to model complex lung diseases that are yet to be modeled and have no cure (e.g., Idiopathic pulmonary fibrosis, etc).

Third, the value of the ALO models is further enhanced due to the availability of companion readouts/ biomarkers (e.g., ViP signatures in the case of respiratory viral pandemics, or monitoring the E gene, or viral shedding, etc.) that can rapidly and objectively vet treatment efficacy based on set therapeutic goals. Of these readouts, the host response, as assessed by ViP signatures, is a key vantage point because an overzealous host response is what is known to cause fatality. Recently, a systematic review of the existing pre-clinical animal models revealed that most of the animal models of COVID-19 recapitulated mild patterns of human COVID-19; no severe illness associated with mortality was observed, suggesting a wide gap between COVID-19 in humans ^38^ and animal models ^90^. It is noteworthy that alternative models that effectively support viral replication, such as the proximal airway epithelium or iPSC-derived AT2 cells (analyzed in this work) or a fetal lung bud tip-derived organoid model recently described by others^77^, do not recapitulate the host response in COVID-19.The model revealed here, in conjunction with the ViP signatures described earlier ^76^, could serve as a pre-clinical model with companion diagnostics to identify drugs that target both the viral and host response in pandemics.

### Limitations of the study

Our adult stem-cell-derived lung organoids, complete with all epithelial cell types, can model COVID-19, but still remains a simplified/rudimentary version compared to the adult human organ. For instance, although the epithelial contributions to the host response are important, it alone cannot account for the response of the immune cells and the non-immune stromal cells, and their crosstalk with the epithelium. Given that epithelial inflammation and damage is propagated by vicious forward-feedback loops of multicellular crosstalk, it is entirely possible that the epithelial signatures induced in infected ALO-derived monolayers are also only a fraction of the actual epithelial response mounted *in vivo*. Regardless of the missing components, what appears to be the case is that we already have a model that recapitulates a ¼ ^th^ to ½ of the genes that are induced across diverse COVID-19 infected patient samples. This model can be further improved by the simultaneous addition of endothelial cells and immune cells to better understand the pathophysiologic basis for DAD, microangiopathy, and even organizing fibrosis with loss of lung capacity that has been observed in many patients^38^; these insights should be valuable to fight some of the long-term sequelae of COVID-19. Future work with flow cytometry and cell sorting of our lung organoids would help understand each cell type’s role in viral pathogenesis. Larger living biobanks of genotyped and phenotyped ALOs, representing donors of different age, ethnicity, predisposing conditions and co-existing comorbidities, will advance our understanding of why SARS-CoV-2 and possibly other infectious agents may trigger different disease course in different hosts. Although we provide proof-of-concept studies in low throughput mode demonstrating the usefulness of the ALOs as human pre-clinical models for screening therapeutics in Phase ‘0’ trials, optimization for the same to be adapted in HTP mode was not attempted here.

## Acknowledgements

This work was supported by the National Institutes for Health (NIH) grants 1R01DK107585-01A1, 3R01DK107585-05S1 (to SD); R01-AI141630, CA100768 and CA160911 (to PG) and R01-AI 155696 (to PG, DS and SD); R00-CA151673 and R01-GM138385 (to DS), R01- HL32225 (to PT), UCOP-R00RG2642 (to SD and PG), UCOP-R01RG3780 (to P.G. and D.S) and a pilot award from the Sanford Stem Cell Clinical Center at UC San Diego Health (P.G, S.D, D.S). GDK was supported through The American Association of Immunologists Intersect Fellowship Program for Computational Scientists and Immunologists. L.C.A’s salary was supported in part by the VA San Diego Healthcare System. The authors would like to thank Victor Pretorius, Rachel White and Jen Bigbee (Department of Cardiothoracic Surgery, UC San Diego) who assisted with thoracotomies during rapid autopsies. This manuscript includes data generated at the UC San Diego Institute of Genomic Medicine (IGC) using an Illumina NovaSeq 6000 that was purchased with funding from a National Institutes of Health SIG grant (#S10 OD026929).

## Author contributions

C.T, M.F, A.F, S.D, P.G and D.S conceptualized the study, C.T, M.F, A.F, S.T, SR.I, N.B, G.D.K, A.C, V.C, M.H, H.R, J.D, L.C.A, A.T, G.L, P.A.T, R.C, T.F.R, D.S., S.D., and P.G. participated in investigation, data curation and formal analysis. C.T, M.F, A.F conducted all experiments on adult lung-derived 3D-organoids and 2D- monolayers; M.F, R.C conducted experiments on iPSC-derived AT2 cells; M.F and G.D.K conducted the flow cytometry analyses; S.T, D.S, P.G conducted the computational analyses; D.S., P.G., S.D. S.T., M.F., A.F, C.T carried out data curation and analyses; A.C., V.C., J.D., L.C.A carried out the rapid autopsies on deceased COVID-19 patients; T.F.R, N.B carried out the studies requiring SARS-CoV-2 infection; J.D., L.C.A, A.T, H.R, P.A.T, M.H provided access to patient tissues. D.S., S.D., P.G. acquired funding for this work; D.S provided the software for computational analyses; P.G. and S.D. drafted the original and revised drafts.

## Data availability

All RNA sequencing data generated by us was deposited in Gene Expression Omnibus (NCBI GEO database) under accession no GSE157055 and GSE157057.

## Key Resource Table

Key Resource Table:
| MATERIALS & REAGENTS |  |  |  |
| --- | --- | --- | --- |
| ANTIBODIES USED FOR IMMUNOFLUORESCENCE |  |  |  |
| Name | Manufacturer | Catalog number | Dilution factor |
| ACE2 | Santa Cruz | 390851 | 1:100 |
| ACE2 | Biolegend | 375802 | 1:50 |
| Acetylated α-Tubulin | Santa Cruz | 23950 | 1:500 |
| AQP5 | Santa Cruz | 514022 | 1:100 |
| CC10 | Santa Cruz | 365992 | 1:100 |
| DAPI | Invitrogen | D1306 | 1:500 |
| KRT5 | Abcam | 52635 | 1:100 |
| MUC5AC | Abcam | 229451 | 1:150 |
| Na<sup>+</sup>/K<sup>+</sup> ATPase | Abcam | 76020 | 1:400 |
| SARS-CoV Nucleoprotein | Sino Biological | 40143-MM05 | 1:250 |
| SFTPB | Santa Cruz | 133143 | 1:100 |
| SFTPC | Abcam | 90716 | 1:150 |
| Occludin | Thermo Fisher | OC-3F10 | 1:500 |
| anti-Rat IgG Secondary Antibody, Alexa Fluor 594 | Invitrogen | A-11007 | 1:500 |
| anti-Rabbit IgG Secondary Antibody, Alexa Fluor 594 | Invitrogen | A-11012 | 1:500 |
| anti-Mouse IgG Secondary Antibody, Alexa Fluor 488 | Invitrogen | A-11011 | 1:500 |
| ANTIBODIES USED FOR IHC |  |  |  |
| SARS Spike (SARS-CoV2) | Abcam | 273433 | 1:250 |
| SARS-CoV Nucleoprotein | Sino Biological | 40143-MM05 | 1:500 |
| ANTIBODIES USED FOR IHC |  |  |  |
| Acetylated-α-tubulin | Santa Cruz | 23950 | 1:8000 |
| AQP5 | Santa Cruz | 514022 | 1:800 |
| KRT5 | Abcam | ab52635 | 1:8000 |
| MUC5 | Abcam | ab229451 | 1:800 |
| SFTPB | Santa Cruz | 133143 | 1:8000 |
| Propidium Iodide | Invitrogen | V13241 | 1:100 |
| anti-Mouse IgG Secondary Antibody, Alexa Fluor 488 | Invitrogen | A-11001 | 1:1000 |
| anti-Rabbit IgG Secondary Antibody, Alexa Fluor 488 | Abcam | ab150077 | 1:1000 |

| INSTRUMENTS |  |  |  |
| --- | --- | --- | --- |
| Countess II Automated Cell Counter | Thermo Fisher Scientific | AMQAX1000 |  |
| Epithelial Volt-Ohm (TEER) Meter | Millipore | MERS00002 |  |
| Leica DMI4000B (Automated Inverted Microscope) | Leica Microsystems | DMI4000B |  |
| Leica TCS SPE Confocal | Leica Microsystems | TCS SPE |  |
| Power Pressure Cooker XL | Tristar Products |  |  |
| Canon Rebel XS DLSR | Canon |  |  |
| MiniAmp Plus Thermal Cycler | Applied Biosystems | A37835 |  |
| QuantStudio5 | Applied Biosystems | A28140 |  |
| Light Microscope (brightfield images) | Carl Zeiss LLC | Axio Observer, Inverted; 491917-0001-000 |  |
| Fisherbrand™ 150 Handheld Homogenizer | Fisher Scientific | 15340168 |  |
| Spark 20M Multimode Microplate Reader | Tecan |  |  |
| Guava® easyCyte Benchtop Flow Cytometer | Millipore | Guava easyCyte 6 2L |  |
| SOFTWARE |  |  |  |
| ImageJ | https://imagej.nih.gov/ij/index.html |  |  |
| GraphPad Prism | https://www.graphpad.com/scientific-software/prism/ |  |  |
| LAS AF Software | https://www.leica-microsystems.com/products/microscope-software/p/leica-las-x-ls/ |  |  |
| QuantStudio Design & Analysis Software | https://www.thermofisher.com/us/en/home/global/forms/life-science/quantstudio-3-5-software.html |  |  |
| CIBERSORTx | https://cibersortx.stanford.edu/runcibersortx.php |  |  |
| FlowJo | Flow Jo V10, BD BioSciences |  |  |
| ENZYMES, CHEMICALS, AND REAGENTS |  |  |  |
| Zinc Formalin | Fisher Scientific | 23-313096 |  |
| Xylene | VWR | XX0060-4 |  |
| Hematoxylin | Sigma-Aldrich Inc | MHS1 |  |
| Ethanol | Koptec | UN1170 |  |
| Sodium Citrate | Sigma-Aldrich | W302600 |  |
| DAB (10x) | Thermo Fisher | 1855920 | 1:10 |
| Stable Peroxidase substrate buffer (10x) | Thermo Fisher | 34062 | 1:10 |
| 3% Hydrogen Peroxide | Target | 245-07-3628 |  |
| Horse Serum | Vector Labs | 30022 |  |
| Anti- rabbit IgG | Vector Laboratories | MP-7401 |  |
| Paraformaldehyde 16% Solution, EM Grade | Electron Microscopy Sciences | 15710 |  |
| 100% Methanol | Supelco | MX0485 |  |
| Glycine | Fisher Scientific | BP381-5 |  |
| **Bovine Serum Albumin** | Sigma-Aldrich | A9647-100G |  |
| **Triton-X 100** | Sigma-Aldrich | X100-500ML |  |
| **Prolong Glass** | Invitrogen | P36984 |  |
| **Nail Polish (Rapid Dry)** | Electron Microscopy Sciences | 72180 |  |
| **Gill Modified Hematoxylin (Solution II)** | Millipore Sigma | 65066-85 |  |
| **Histogel** | Thermo Scientific | HG4000012 |  |
| **TrypLE Select** | Thermo Scientific | 12563-011 |  |
| **Advanced DMEM/F-12** | Thermo Scientific | 12634-010 |  |
| **HEPES Buffer** | Life Technologies | 15630080 |  |
| **Glutamax** | Thermo Scientific | 35050-061 |  |
| **Penicillin-Streptomycin** | Thermo Scientific | 15140-122 |  |
| **Collagenase Type I** | Thermo Scientific | 17100-017 |  |
| **Matrigel** | Corning | 354234 |  |
| **B-27** | Thermo Scientific | 17504044 |  |
| **N-acetyl-L-cysteine** | Sigma-Aldrich | A9165 |  |
| **Nicotinamide** | Sigma-Aldrich | N0636 |  |
| **FGF-7 (KGF)** | PeproTech | 100-19-50ug |  |
| **FGF10** | PeproTech | 100-26-50ug |  |
| **A-83-01** | Bio-Techne Sales Corp. | 2939/50 |  |
| **SB202190** | Sigma-Aldrich | S7067-25MG |  |
| **Y-27632** | R&D Systems | 1254/50 |  |
| **DPBS** | Thermo Scientific | 14190-144 |  |
| **Ultrapure Water** | Invitrogen | 10977-015 |  |
| **EDTA** | Thermo Scientific | AM9260G |  |
| **Hydrocortisone** | STEMCELL Technologies | 7925 |  |
| **Heparin** | Sigma Aldrich | H3149 |  |
| **Pneumacult Ex-Plus Medium** | STEMCELL Technologies | 5040 |  |
| **PneumaCult ALI Medium** | STEMCELL Technologies | 5001 |  |
| **Goat serum** | Vector Laboratories | MP-7401 |  |
| **Fetal Bovine Serum** | Sigma-Aldrich | F2442-500ML |  |
| **Animal Component-Free Cell Dissociation Kit** | STEMCELL Technologies | 5426 |  |
| **Red Blood Cell Lysis Buffer** | Invitrogen | 00-4333-57 |  |
| **Cell Recovery Solution** | Corning | 354253 |  |
| **Sodium Azide** | Fisher Scientific | S227I-100 |  |
| **Cyto-Fast Fix/Perm Buffer Set** | Biolegend | 426803 |  |
| **FITC-Dextran** | Sigma-Aldrich | FD10S |  |
| **Quick-RNA MicroPrep Kit** | Zymo Research | R1051 |  |
| **Quick-RNA MiniPrep Kit** | Zymo Research | R1054 |  |
| **Ethyl alcohol, pure** | Sigma-Aldrich | E7023 |  |
| **TRI Reagent** | Zymo Research | R2050-1-200 |  |
| **2x SYBR Green qPCR Master Mix** | Bimake | B21203 |  |
| **qScript cDNA SuperMix** | Quanta Biosciences | 95048 |  |
| **Applied Biosystems TaqMan Fast Advanced Master Mix** | Thermo Scientific | 4444557 |  |
| **18S, Hs99999901_s1** | Thermo Scientific | 4331182 |  |
| **E_Sarbeco_F1 Forward Primer** | IDT | 10006888 |  |
| **E_Sarbeco_R2 Reverse Primer** | IDT | 10006890 |  |
| **E_Sarbeco_P1 Probe** | IDT | 10006892 |  |
| **OTHER** |  |  |  |
| **6-well Tissue Culture Plate** | Genesee Scientific | 25-105 |  |
| **12-well Tissue Culture Plate** | CytoOne | CC7682-7512 |  |
| **Transwell Inserts (6.5 mm, 0.4 um pore size)** | Corning | 3470 |  |
| **Cell Scraper** | Millipore Sigma | C5981-100EA |  |
| **Microscope Cover Glass (#1 Thickness) 24 x 50 mm** | VWR | 16004-098 |  |
| **Microscope Cover Glass (#1 Thickness) 25 mm diameter** | Chemglass Life Sciences | CLS-1760-025 |  |
| **Millicell EZ Slide 8-Well Chamber** | Millipore Sigma | PEZGS0816 |  |
| **Optical Lens Cleaner** | Fisher Scientific | 22-143974 |  |
| **Countess Cell Counting Chamber Slides** | Invitrogen | C10312 |  |
| **Trypan Blue Stain** | Invitrogen | T10282 |  |
| **70 um Cell Strainer** | Thermo Fisher Scientific | 22-363-548 |  |
| **100 um Cell Strainer** | Corning | 352360 |  |
| **RNase Away** | Thermo Fisher Scientific | 14-375-35 |  |
| **Noyes Spring Scissors - Angled** | Fine Science Tools | 15013-12 |  |

## Detailed Methods

### Collection of human lung specimens for organoid isolation

To generate adult healthy lung organoids, fresh biopsy bites were prospectively collected after surgical resection of the lung by the cardiothoracic surgeon. Before collection of the lung specimens, each tissue was sent to a gross anatomy room where a pathologist cataloged the area of focus, and the extra specimens were routed to the research lab in Human Transport Media (HTM, Advanced DMEM/F-12, 10 mM HEPES, 1X Glutamax, 1X penicillin-streptomycin, 5 μM Y-27632) for cell isolation. Deidentified lung tissues obtained during surgical resection, that were deemed excess by clinical pathologists, were collected using an approved human research protocol (IRB# 101590; PI: Thistlethwaite). Isolation and biobanking of organoids from these lung tissues were carried out using an approved human research protocol (IRB# 190105: PI Ghosh and Das) that covers human subject research at the UC San Diego HUMANOID Center of Research Excellence (CoRE). For all the deidentified human subjects, information including age, gender, and previous history of the disease, was collected from the chart following the rules of HIPAA and described in the Table.

A portion of the same lung tissue specimen was fixed in 10% Zinc-Formalin for at least 24hrs followed by submersion in 70% EtOH until embedding in FFPE blocks.

### Autopsy procedures for lung tissue collection from COVID-19 positive human subjects

The lung specimens from COVID-19 positive human subjects were collected through autopsy (the study was IRB Exempt). All donations to this trial were obtained after telephone consent followed by written email confirmation by the next of kin/power of attorney per California state law (no in-person visitation could be allowed into our COVID-19 ICU during the pandemic). The team member followed the CDC guidelines for COVID19 and the autopsy procedures^91, 92^). Lung specimens were collected in 10% Zinc-formalin and stored for 72 hrs before processing for histology. Patient characteristics are listed in the Table.

Autopsy #2 was a standard autopsy performed by anatomical pathology in the BSL3 autopsy suite. The patient expired and his family consented for autopsy. After 48 hrs, the lungs were removed and immersion fixed whole in 10% formalin for 48 hrs and then processed further. Lungs were only partially fixed at this time (about 50% fixed in thicker segments) and were sectioned into small 2-4 cm chunks and immersed in 10% formalin for further investigation.

Autopsy #4 and #5 were collected from rapid post-mortem lung biopsies. The procedure was performed in the Jacobs Medical Center ICU (all of the ICU rooms have a pressure-negative environment, with air exhausted through HEPA filters [Biosafety Level 3 (BSL3)] for isolation of SARS-CoV-2 virus). Biopsies were performed 2-4 hrs after patient expiration. The ventilator was shut off to reduce the aerosolization of viral particles at least 1 hr after the loss of pulse and before sample collection. Every team member had personal protective equipment in accordance with the University policies for procedures on patients with COVID-19 (N95 mask + surgical mask, hairnet, full face shield, surgical gowns, double surgical gloves, booties). Lung biopsies were obtained after L-thoracotomy in the 5th intercostal space by the cardiothoracic surgery team. Samples were taken from the left upper lobe (LUL) and left lower lobe (LLL) and then sectioned further.

### Isolation and culture of human whole lung-derived organoids

A previously published protocol was modified to isolate lung organoids from 3 human subjects ^31, 33^. Briefly, normal human lung specimens were washed with PBS/4X penicillin-streptomycin and minced with surgical scissors. Tissue fragments were resuspended in 10 mL of wash buffer (Advanced DMEM/F-12, 10 mM HEPES, 1X Glutamax, 1X penicillin-streptomycin) containing 2 mg/ml Collagenase Type I (Thermo Fisher, USA) and incubated at 37°C for approximately 1 hr. During incubation, tissue pieces were sheared every 10 min with a 10 mL serological pipette and examined under a light microscope to monitor the progress of digestion. When 80-100% of single cells were released from connective tissue, the digestion buffer was neutralized with 10 mL wash buffer with added 2% Fetal Bovine Serum; the suspension was passed through a 100-µm cell strainer and centrifuged at 200 rcf. Remaining erythrocytes were lysed in 2 ml red blood cell lysis buffer (Invitrogen) at room temperature for 5 min, followed by the addition of 10 mL of wash buffer and centrifugation at 200 rcf. Cell pellets were resuspended in cold Matrigel (Corning, USA) and seeded in 25 µl droplets on a 12 well tissue culture plate. The plate was inverted and incubated at 37°C for 10 min to allow complete polymerization of the Matrigel before the addition of 1 mL Lung Expansion Medium per well. Lung expansion media was prepared by modifying a media that was optimized previously for growing gastrointestinal (GI)-organoids (50% conditioned media, prepared from L-WRN cells with Wnt3a, R-spondin, and Noggin, ATCC-CRL-3276^TM^) ^42, 93–95^ with a proprietary cocktail from the HUMANOID CoRE containing B27, TGF-β receptor inhibitor, antioxidants, p38 MAPK inhibitor, FGF 7, FGF 10 and ROCK inhibitor. The lung expansion media was compared to alveolosphere media I (IMDM and F12 as the basal medium with B27, low concentration of KGF, Monothioglycerol, GSK3 inhibitor, Ascorbic acid, Dexamethasone, IBMX, cAMP and ROCK inhibitor) and II (F12 as the basal medium with added CaCl_2_, B27, low concentration of KGF, GSK3 inhibitor, TGF-β receptor inhibitor Dexamethasone, IBMX, cAMP and ROCK inhibitor) modified from previously published literature ^27, 28^. Neither alvelosphere media contain any added Wnt3a, R-spondin, and Noggin. The composition of these media was developed either by fundamentals of adult-stem cell-derived mixed epithelial cellularity in other organs (like the gastrointestinal tract ^40–42^, or rationalized based on published growth conditions for proximal and distal airway components^25, 31, 32^. Organoids were maintained in a humidified incubator at 37°C/5% CO2, with a complete media change performed every 3 days. After the organoids reached confluency between 7-10 days, organoids were collected in PBS/0.5 mM EDTA and centrifuged at 200 rcf for 5 min. Organoids were dissociated in 1 mL trypLE Select (Gibco, USA) per well at 37°C for 4-5 min and mechanically sheared. Wash buffer was added at a 1:5, trypLE to wash buffer ratio. The cell suspension was subsequently centrifuged, resuspended in Matrigel, and seeded at a 1:5 ratio. Lung organoids were biobanked and passage 3-8 cells were used for experiments. Subculture was performed every 7-10 days.

### The preparation of lung organoid-derived monolayers

Lung-organoid-derived monolayers were prepared using a modified protocol of GI-organoid-derived monolayers ^42, 93–95^. Briefly, transwell inserts (6.5 mm diameter, 0.4 um pore size, Corning) were coated in Matrigel diluted in cold PBS at a 1:40 ratio and incubated for 1hr at room temperature. Confluent organoids were collected in PBS/EDTA on day 7 and dissociated into single cells in trypLE for 6-7 min at 37°C. Following enzymatic digestion, the cell suspension was mechanically sheared through vigorous pipetting with a 1000 µl pipette and neutralized with wash buffer. The suspension was centrifuged, resuspended in Pneumacult Ex-Plus Medium (StemCell, Canada), and passed through a 70-µm cell strainer. The coating solution was aspirated, and cells were seeded onto the apical membrane at 1.8E5 cells per transwell with 200 µl PneumaCult Ex-Plus media. 700 µl of PneumaCult Ex-Plus was added to the basal chamber. Cells were cultured over the course of 2-4 days. A media change of both the apical and basal chambers was performed every 24 hrs.

### Air-Liquid Interface (ALI) Model of Lung organoids

Organoids were dissociated into single cells and expanded in T-75 flasks in PneumaCult Ex-Plus Medium until confluency was reached. Cells were dissociated in ACF Enzymatic Dissociation Solution (StemCell, Canada) for 6-7 min at 37°C and neutralized in equal volume ACF Enzyme Inhibition Solution (StemCell, Canada). Cells were seeded in the apical chamber of transwells at 3.3E4 cells per transwell in 200 µL of PneumaCult Ex-Plus Medium. 500 µL of PneumaCult Ex-Plus was added to the basal chamber. Media in both chambers was changed every other day until confluency was reached (∼4 days). The media was completely removed from the apical chamber, and media in the basal chamber was replaced with ALI Maintenance Medium (StemCell, Canada). The media in the basal chamber was changed every 2 days. The apical chamber was washed with warm PBS every 5-7 days to remove accumulated mucus. Cells were cultured under ALI conditions for 21+ days until they completed differentiation into a pseudostratified mucociliary epithelium. To assess the integrity of the epithelial barrier, Trans-Epithelial Electrical Resistance (TEER) was measured with an Epithelial Volt-Ohm Meter (Millicell, USA). The media was removed from the basal chamber, and wash media was added to both chambers. Cultures were equilibrated to 37°C before TEER values were measured. Final values were expressed as Ωꞏcm^2^ units and were calculated by multiplying the growth area of the membrane by the raw TEER value.

### The culture of primary airway epithelial cells and iPSC-derived alveolar epithelial cells

Primary normal human bronchial epithelial cells (NHBE) were obtained from Lonza and grown according to instructions. NHBE cells were cultured in T25 cell culture tissue flasks with PneumaCult-Ex Plus media (StemCell, Canada). Cells were seeded at ∼100,000 cells/T25 flask and incubated at 37°C, 5% CO2. Once cells reached 70–80% confluency, they were dissociated using 0.25% Trypsin in dissociation media and plated in 24 well transwells (Corning). Primary human bronchial epithelial cells (HBEpC) and small airway epithelial cells (HSAEpC) were obtained from Cell Applications Inc. The HBEpC and HSAEpC were cultured in human bronchial/tracheal epithelial cell media and small airway epithelial cell media, respectively, following the instructions of Cell Application.

Human iPSC-derived alveolar epithelial type 2 cells (iHAEpC2) were obtained from Cell Applications Inc. and cultured in growth media (i536K-05, Cell Applications Inc.) according to the manufacturer’s instructions. All the primary cells were used within early passages (5-6) to avoid any gradual disintegration of the airway epithelium with columnar epithelial structure and epithelial integrity.

### The infection with SARS-Cov2

Lung organoid-derived monolayers or primary airway epithelial cells either in 384 well plates or in transwells were washed twice with antibiotic-free lung wash media. 1E5 PFU of SARS-CoV-2 strain USA-WA1/2020 (BEI Resources NR-52281) in complete DMEM was added to the apical side of the transwell and allowed to incubate for 24, 48, 72 and 96 hrs at 34°C and 5% CO2. After incubation, the media was removed from the basal side of the transwell. The apical side of the transwells was then washed twice with (antibiotic-free lung wash media) and then twice with PBS. TRIzol™ Reagent (Thermo Fisher 15596026) was added to the well and incubated at 34°C and 5% CO2 for 10 min. The TRIzol™ Reagent was removed and stored at -80 °C for RNA analysis.

### RNA isolation

Organoids and monolayers used for lung cell type studies were lysed using RNA lysis buffer followed by RNA extraction per Zymo Research Quick-RNA MicroPrep Kit instructions. Tissue samples and monolayers in SARS-CoV2 studies were lysed in TRI-Reagent and RNA was extracted using Zymo Research Direct-zol RNA Miniprep.

### Quantitative (q)RT-PCR

Organoid and monolayer cell-type gene expression was measured by qRT-PCR using 2x SYBR Green qPCR Master Mix. cDNA was amplified with gene-specific primer/probe set for the lung cell type markers and qScript cDNA SuperMix (5x). qRT-PCR was performed with the Applied Biosystems QuantStudio 5 Real-Time PCR System. Cycling parameters were as follows: 95 °C for 20 s, followed by 40 cycles of 1 s at 95 °C and 20 s at 60 °C. All samples were assayed in triplicate and eukaryotic 18S ribosomal RNA was used as a reference.

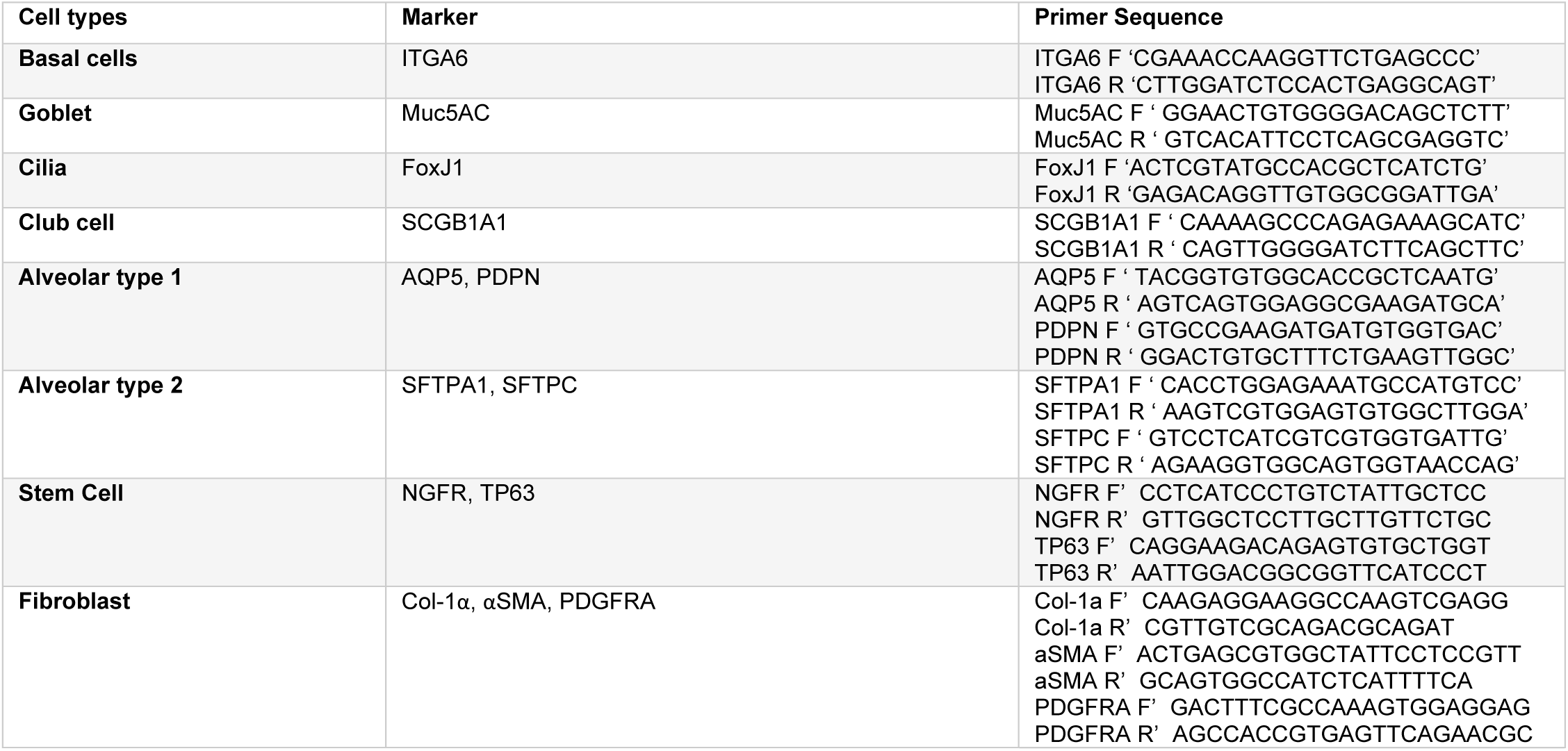

### Assessment of SARS-CoV-2 infectivity test

Assessment of SARS-CoV-2 infectivity test was determined by qPCR using TaqMan assays and TaqMan Universal PCR Master Mix as done before^96, 97^. cDNA was amplified with gene-specific primer/probe set for the E gene and QPCR was performed with the Applied Biosystems QuantStudio 3 Real-Time PCR System. The specific TaqMan primer/probe set for E gene are as follows: Forward 5’-ACAGGTACGTTAATAGTTAATAGCGT-3’, (IDT, Cat# 10006888); Reverse 5’-ATATTGCAGCAGTACGCACACA-3’; Probe 5’-FAM-ACACTAGCCATCCTTACTGCGCTTCG-BBQ-3’ and 18S rRNA. Cycling parameters were as follows: 95 °C for 20 s, followed by 40 cycles of 1 s at 95 °C and 20 s at 60 °C. All samples were assayed in triplicate and gene eukaryotic 18S ribosomal RNA was used as a reference.

### Immunofluorescence

Organoids and Lung organoid-Derived Monolayers were fixed by either: (1) 4% PFA at room temperature for 30 min and quenched with 30 mM glycine for 5 min, (2) ice-cold 100% Methanol at -20°C for 20 min, (3) ice-cold 100% Methanol on ice for 20 min. Subsequently, samples were permeabilized and blocked for 2 hrs using an in-house blocking buffer (2 mg/mL BSA and 0.1% Triton X-100 in PBS); as described previously^98^. Primary antibodies were diluted in blocking buffer and allowed to incubate overnight at 4°C; Secondary antibodies were diluted in blocking buffer and allowed to incubate for 2 hrs in the dark. Antibody dilutions are listed in the Supplementary Key Resource Table. ProLong Glass was used as a mounting medium. #1 Thick Coverslips were applied to slides and sealed. Samples were stored at 4°C until imaged. FFPE embedded Organoid and Lung Tissue sections underwent antigen retrieval as previously described in methods for Immunohistochemistry staining. After antigen retrieval and cooling in DI water; samples were permeabilized and blocked in blocking buffer and treated as mentioned above for immunofluorescence. Images were acquired at room temperature with Leica TCS SPE confocal and with DMI4000 B microscope using the Leica LAS-AF Software. Images were taken with a 40× oil-immersion objective using 405-,488-, 561-nm laser lines for excitation. Z-stack images were acquired by successive Z-slices of 1µm in the desired confocal channels. Fields of view that were representative and/or of interest were determined by randomly imaging 3 different fields. Z-slices of a Z-stack were overlaid to create maximum intensity projection images; all images were processed using FIJI (Image J) software.

### Embedding of Organoids in *HistoGel*

Organoids were seeded on a layer of Matrigel in 6-Well plates and grown for 7-8 days. Once mature, organoids were fixed in 4% PFA at room temperature for 30 min and quenched with 30 mM glycine for 5 min. Organoids were gently washed with PBS and harvested using a cell scraper. Organoids were resuspended in PBS using wide-bore 1000 µL pipette tips. Organoids were stained using Gill’s hematoxylin for 5 min for easier FFPE embedding and sectioning visualization. Hematoxylin stained organoids were gently washed in PBS and centrifuged and excess hematoxylin was aspirated. Organoids were resuspended in 65°C histogel and centrifuged at 65°C for 5 min. Histogel embedded organoid pellets were allowed to cool to room temperature and stored in 70% ethanol at 4°C until ready for FFPE embedding by LJI Histology Core. Successive FFPE embedded organoid sections were cut at a 4 µm thickness and fixed on to microscope slides.

### Immunohistochemistry

For SARS CoV- nucleoprotein (np) antigen retrieval, slides were immersed in Tris-EDTA buffer (pH 9.0) and boiled for 10 min at 100°C inside a pressure cooker. Endogenous peroxidase activity was blocked by incubation with 3% H_2_O_2_ for 10 minutes. To block non-specific protein binding 2.5% goat serum was added. Tissues were then incubated with a rabbit SARS CoV-NP antibody (Sino Biological, See Supplementary Key Resource Table) for 1.5 hrs at room temperature in a humidified chamber and then rinsed with TBS or PBS 3 times, for 5 min each. Sections were incubated with horse anti-rabbit IgG secondary antibodies for 30 min at room temperature and then washed with TBS or PBS 3 times for 5 min each. Sections were incubated with DAB and counterstained with hematoxylin for 30 sec.

### Permeability of lung monolayer using FITC-Dextran

Adult lung monolayers were grown for 2 days in PneumaCult Ex-Plus media on transwell inserts (6.5 mm diameter, 0.4 um pore size, Corning). Trans-Epithelial Electrical Resistance (TEER) was monitored with an Epithelial Volt-Ohm Meter (Millicell, USA). On the second day of growth, FITC-dextran (10 kD) was added at a 1:50 dilution in lung wash media. The basolateral side of the insert was changed to lung wash media only. After 30 minutes of incubation with FITC-dextran, 50 µl of the basolateral supernatant was transferred to an opaque welled 96-well plate. Fluorescence was measured using a TECAN plate reader.

### The characterization of lung cell types using Flow Cytometry

Lung organoids were dissociated into single cells via trypLE digestion and strained with a 30 µm filter (Miltenyi Biotec, Germany). Approximately 2.5E5 cells for each sample were fixed and permeabilized at room temperature in Cyto-Fast Fix Perm buffer (BioLegend, USA) for 20 min. The samples were subsequently washed with Cyto-Fast Perm Wash solution (BioLegend, USA) and incubated with lung epithelial cell type markers for 30 min. Following primary antibody incubation, the samples were washed and incubated with Propidium Iodide (Invitrogen) and Alexa Flour 488 secondary antibodies (Invitrogen) for 30 min. Samples were resuspended in FACS buffer (PBS, 5% FBS, 2 mM Sodium Azide). Flow cytometry was performed using Guava® easyCyte benchtop flow cytometer (Millipore) and data was analyzed using InCyte (version 3.3) and FlowJo X v10 software.

### RNA Sequencing

RNA sequencing libraries were generated using the Illumina TruSeq Stranded Total RNA Library Prep Gold with TruSeq Unique Dual Indexes (Illumina, San Diego, CA). Samples were processed following manufacturer’s instructions, except modifying RNA shear time to five minutes. The resulting libraries were multiplexed and sequenced with 100 basepair (bp) Paired-End (PE100) to a depth of approximately 25-40 million reads per sample on an Illumina NovaSeq 6000 by the Institute of Genomic Medicine (IGM) at the University of California San Diego. Samples were demultiplexed using bcl2fastq v2.20 Conversion Software (Illumina, San Diego, CA). RNASeq data was processed using kallisto (version 0.45.0), and human genome GRCh38 Ensembl version 94 annotation (Homo_sapiens GRCh38.94 chr_patch_hapl_scaff.gtf). Gene-level TPM values and gene annotations were computed using tximport and biomaRt R package. A custom python script was used to organize the data and log reduced using log2(TPM+1). The raw data and processed data are deposited in Gene Expression Omnibus under accession no GSE157055, and GSE157057.

### Data Collection and Annotation

Publicly available COVID-19 gene expression databases were downloaded from the National Center for Biotechnology Information (NCBI) Gene Expression Omnibus website (GEO) ^99–101^. If the dataset is not normalized, RMA (Robust Multichip Average)^102, 103^ is used for microarrays and TPM (Transcripts Per Millions)^104, 105^ is used for RNASeq data for normalization. We used log2(TPM+1) to compute the final log-reduced expression values for RNASeq data. Accession numbers for these crowdsourced datasets are provided in the figures and manuscript. All of the above datasets were processed using the Hegemon data analysis framework^106–108^.

### Analysis of RNA seq Datasets

DESeq2 ^109^ was applied to uninfected and infected samples to identify Up- and Down-regulated genes. A gene signature score is computed using both the Up- and Down-regulated genes which are used to order the sample. To compute the gene signature score, first, the genes present in this list were normalized according to a modified Z-score approach centered around StepMiner threshold (formula = (expr -SThr)/3*stddev). The normalized expression values for every probeset for all the genes were added or subtracted (depending on Up and Down-regulated genes) together to create the final score. The samples were ordered based on the final gene signature score. The Gene signature score is used to classify sample categories and the performance of the multi-class classification is measured by ROC-AUC (Receiver Operating Characteristics Area Under The Curve) values. A color-coded bar plot is combined with a violin plot to visualize the gene signature-based classification. All statistical tests were performed using R version 3.2.3 (2015-12-10). Standard t-tests were performed using python scipy.stats.ttest_ind package (version 0.19.0) with Welch’s Two Sample t-test (unpaired, unequal variance (equal_var=False), and unequal sample size) parameters. Multiple hypothesis correction was performed by adjusting *p* values with statsmodels.stats.multitest.multipletests (fdr_bh: Benjamini/Hochberg principles). The results were independently validated with R statistical software (R version 3.6.1; 2019-07-05). Pathway analysis of gene lists was carried out via the Reactome database and algorithm ^110^. Reactome identifies signaling and metabolic molecules and organizes their relations into biological pathways and processes. Violin, Swarm and Bubble plots are created using python seaborn package version 0.10.1.

### Single Cell RNA Seq data analysis

Single Cell RNASeq data from GSE145926 was downloaded from GEO in the HDF5 Feature Barcode Matrix Format. The filtered barcode data matrix was processed using Seurat v3 R package^111^ and scanpy python package^112^. Pseudo bulk analysis of GSE145926 data was performed by adding counts from the different cell subtypes and normalized using log2(CPM+1). Epithelial cells were identified using SFTPA1, SFTPB, AGER, AQP4, SFTPC, SCGB3A2, KRT5, CYP2F1, CCDC153, and TPPP3 genes using SCINA algorithm^113^. Pseudo bulk datasets were prepared by adding counts from the selected cells and normalized using log (CPM+1).

### Assessment of cell-type proportions

CIBERSORTx (https://cibersortx.stanford.edu/runcibersortx.php) was used for cell-type deconvolution of our dataset (which was normalized by CPM). As reference samples, we first used the Single Cell RNASeq dataset (GSE132914) from Gene Expression Omnibus (GEO). Next, we analyzed the bulk RNA seq datasets for the identification of cell types of interest using relevant gene markers (see **Table 2**): AT1 cells (PDPN, AQP5, P2RX4, TIMP3, SERPINE1), AT2 cells (SFTPA1, SFTPB, SFTPC, SFTPD, SCGB1A1, ABCA3, LAMP3), BASAL cells (CD44, KRT5, KRT13, KRT14, CKAP4, NGFR, ITGA6), CLUB cells (SCGB1A1, SCGB3A2, SFTPA1, SFTPB, SFTPD, ITGA6, CYP2F1), GOBLET cells (CDX2, MUC5AC, MUC5B, TFF3), Ciliated cells (ACTG2, TUBB4A, FOXA3, FOXJ1, SNTN), Generic Lung Lineage cells (GJA1, TTF1, EPCAM) were identified using SCINA algorithm. Then, normalized pseudo counts were obtained with the CPM normalization method. The cell-type signature matrix was derived from the Single Cell RNASeq dataset, cell-types, and gene markers of interest. It was constructed by taking an average from gene expression levels for each of the markers across each cell type.

### Statistical analysis

All experiments were repeated at least three times, and results were presented either as one representative experiment or as average ± S.E.M. Statistical significance between datasets with three or more experimental groups was determined using one-way ANOVA including a Tukey’s test for multiple comparisons. For all tests, a P-value of 0.05 was used as the cutoff to determine significance (*P < 0.05, **P < 0.01, ***P < 0.001, and ****P < 0.0001). All experiments were repeated a least three times, and P-values are indicated in each figure. All statistical analyses were performed using GraphPad prism 6.1. A part of the statistical tests was performed using R version 3.2.3 (2015-12-10). Standard t-tests were performed using python scipy.stats.ttest_ind package (version 0.19.0).

---

**Figure 1- figure supplement 1.**
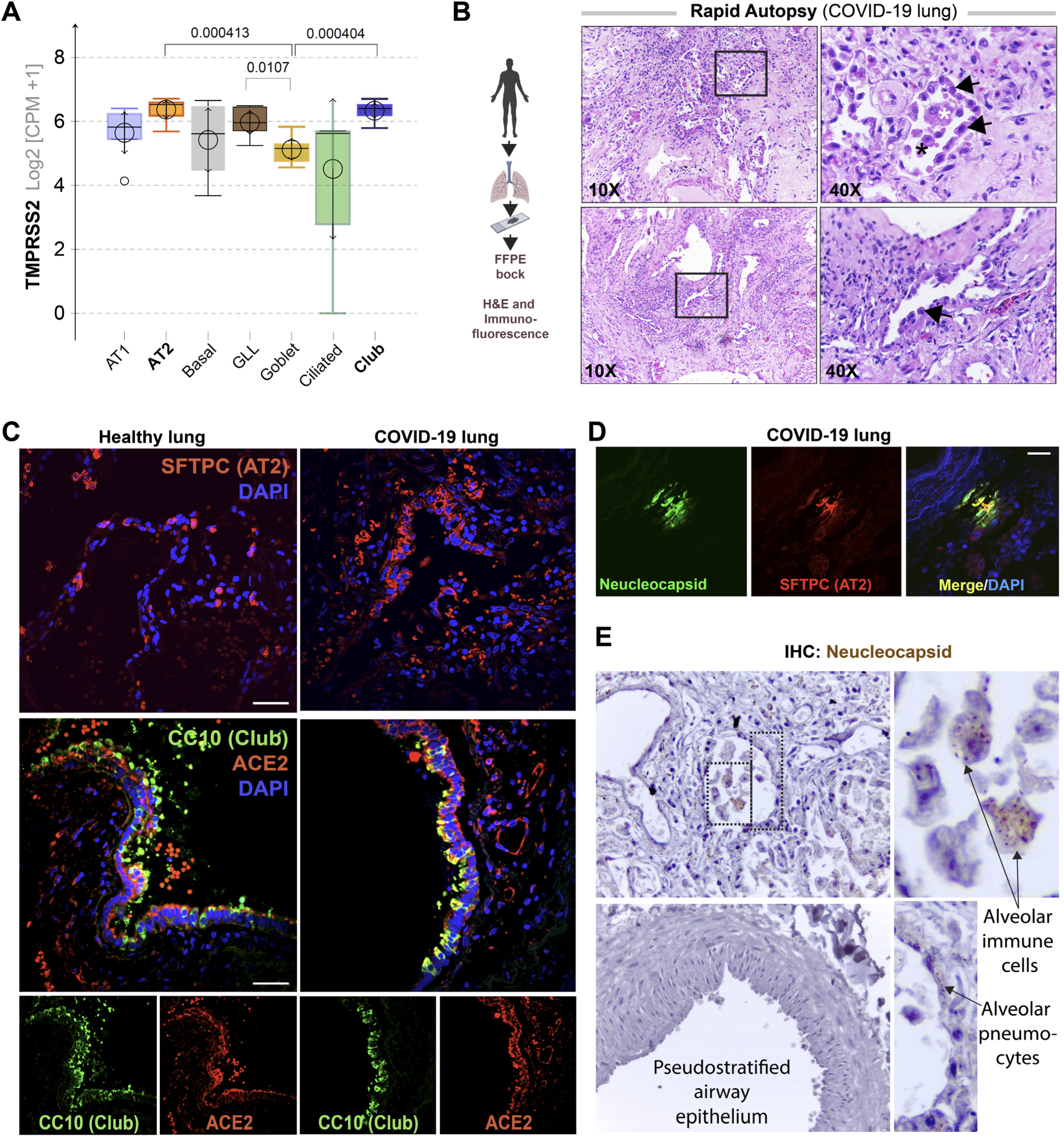
**Alveolar type II pneumocyte hyperplasia is a pathognomonic feature of lung injury in COVID-19.** **A.**Whisker plots display relative levels of TMPRSS2 expression in various cell types in the normal human lung. The cell types were annotated within a publicly available single-cell sequencing dataset (GSE132914) using genes listed in Table 2. *p*-values were analyzed by one-way Anova and Tukey post hoc test. **B.** FFPE sections of the human lung from deceased COVID-19 patients were analyzed by H&E staining. Representative fields are shown. Images on the right are magnified areas indicated with boxes on the left. Arrows = alveolar type II pneumocyte hyperplasia. **C-D.** FFPE sections of the human lung from normal and deceased COVID-19 patients were stained for AT2 and club cell markers and either ACE2 or viral nucleocapsid protein and analyzed by confocal immunofluorescence. Representative images are shown. Scale bar = 50 µm. **E**. FFPE sections of the human lung from normal and deceased COVID-19 patients were stained for viral nucleocapsid antibody. Representative images are shown. Arrows = infected cells.

**Figure 2- Figure Supplement 1.**
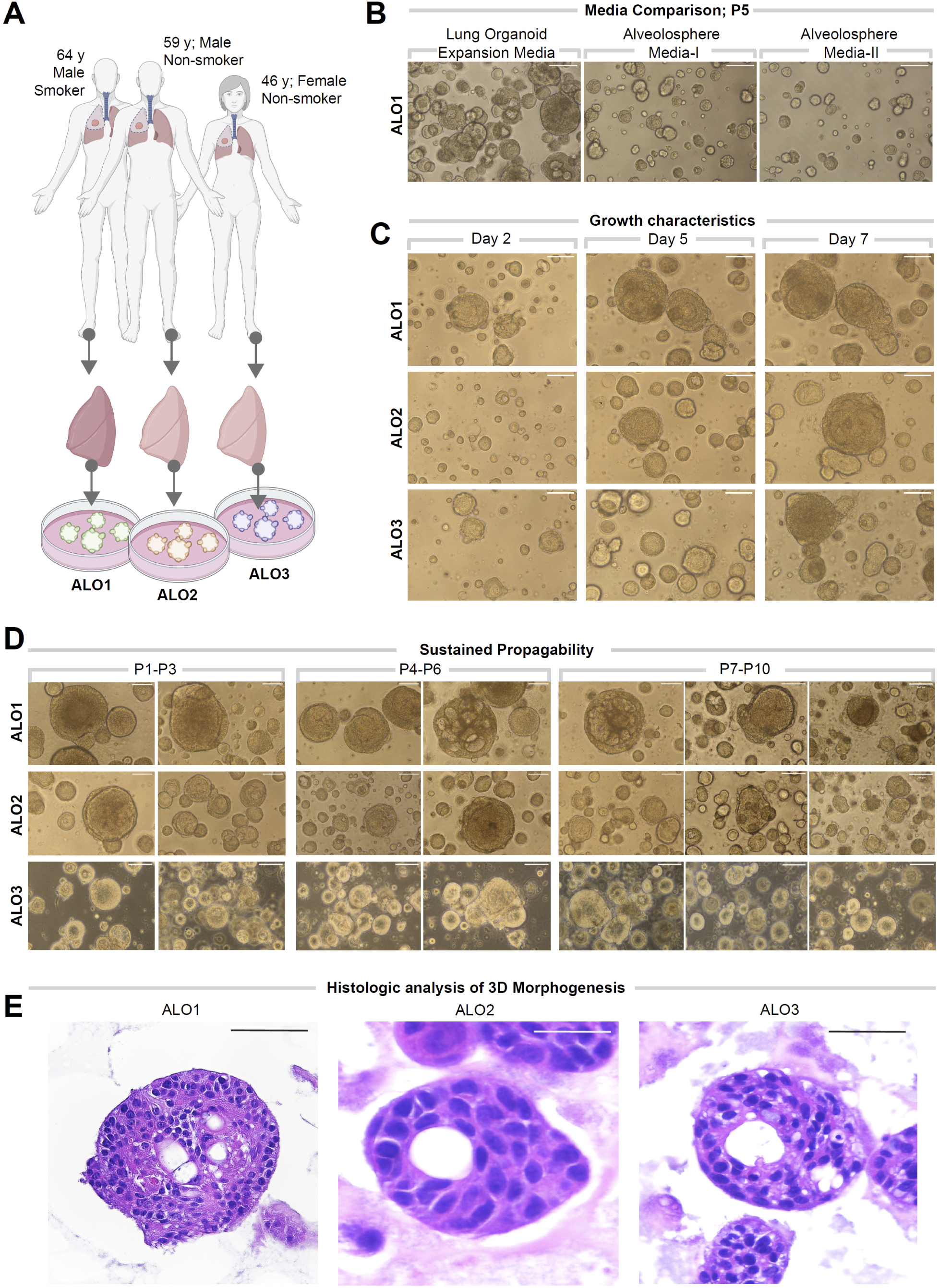
**Lung organoids are reproducibly established from 3 different donors and propagated in each case over 10 passages.** **A.**Schematic displaying the key demographics of the patients who served as donors of the lung tissue as a source of adult stem cells for the generation of organoids. Three organoid lines were generated, ALO1-3. ALO, adult lung organoids. **B-D**. Bright-field microscopy of organoids in 3D culture grown in different media/conditions (B), imaged serially over days (C) and at different passages (D). Scale bar = 100 µm. **E**. Serial cuts of *HistoGel-*embedded organoids were analyzed by H&E staining. Scale bar = 50 µm.

**Figure 2- Figure Supplement 2.**
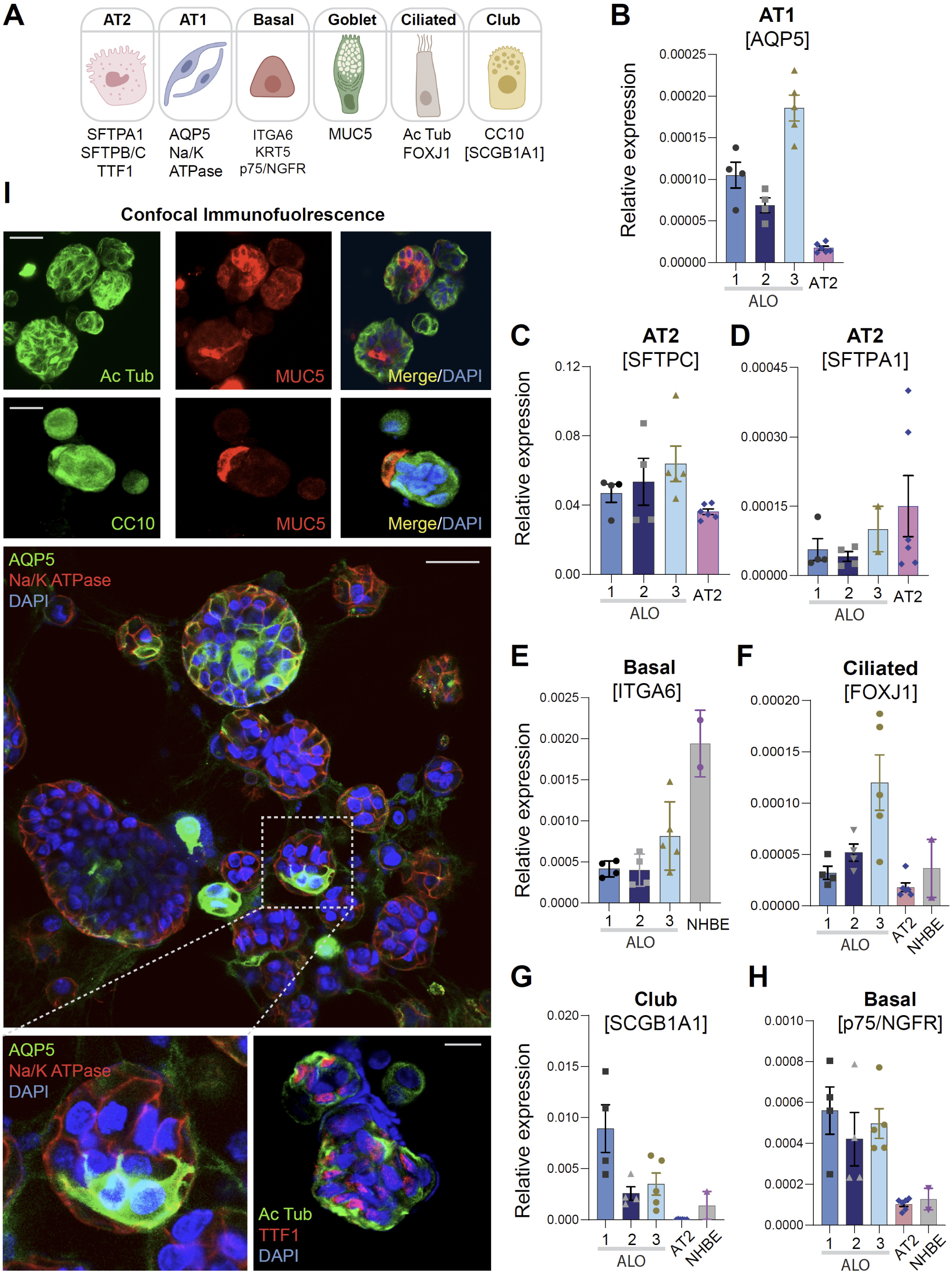
**Adult stem cell-derived lung organoids are propagatable models with both proximal and distal airway components.** **A.**Schematic lists the various markers used here for qPCR and immunofluorescence to confirm the presence of all cell types in the 3D lung organoids here and in 2D monolayers later (in Fig 3). **B-H.** Bar graphs display the relative abundance of various cell type markers (normalized to 18S) in adult lung organoids (ALO), compared to the airway (NHBE) and/or alveolar (AT2) control cells, as appropriate. *p*-values were analyzed by one-way Anova. Error bars denote S.E.M; n = 3-6 datasets. **I.** 3D organoids grown in 8-well chamber slides were fixed, immunostained and visualized by confocal microscopy, as in Fig 2K. Scale bar = 50 µm.

**Figure 2- Figure Supplement 3.**
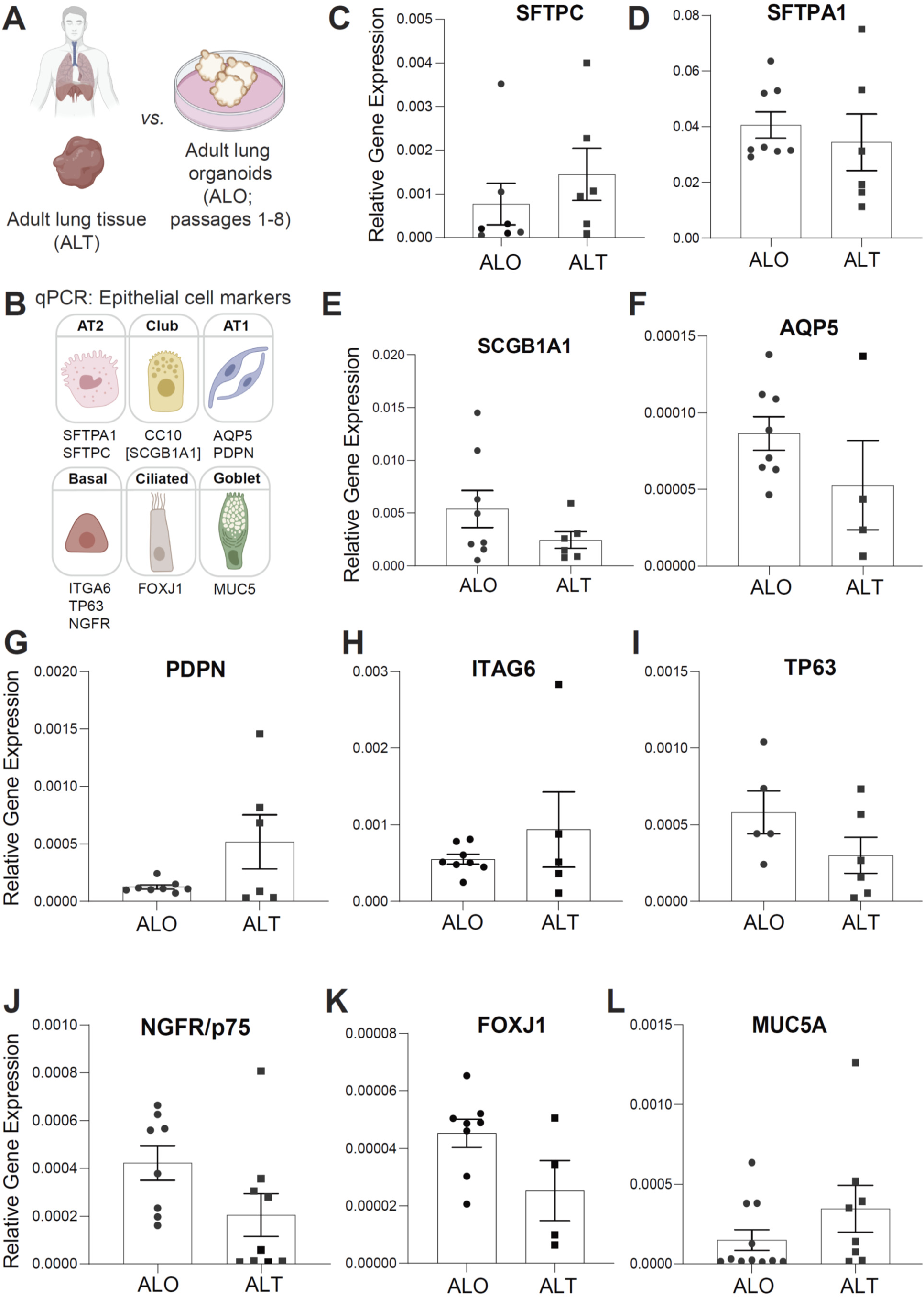
**Adult stem cell-derived lung organoids (ALO) generally recapitulate cell type specific gene expression patterns observed in the adult lung tissue (ALT) from which they originate.** **A-B.** Schematics depict the study goal in this figure, i.e., analysis of cell type specific transcripts in ALO vs ALT. **C-L.** Bar graphs display the relative abundance of various cell type markers (normalized to 18S) in adult lung organoids from early passage (ALO), compared to the adult lung tissue (ALT) from which they were derived. *p*-values were analyzed by one-way Anova. Error bars denote S.E.M; n = 3-6 datasets. Statistically significant differences were not noted in any of the transcripts analyzed.

**Figure 2- Figure Supplement 4.**
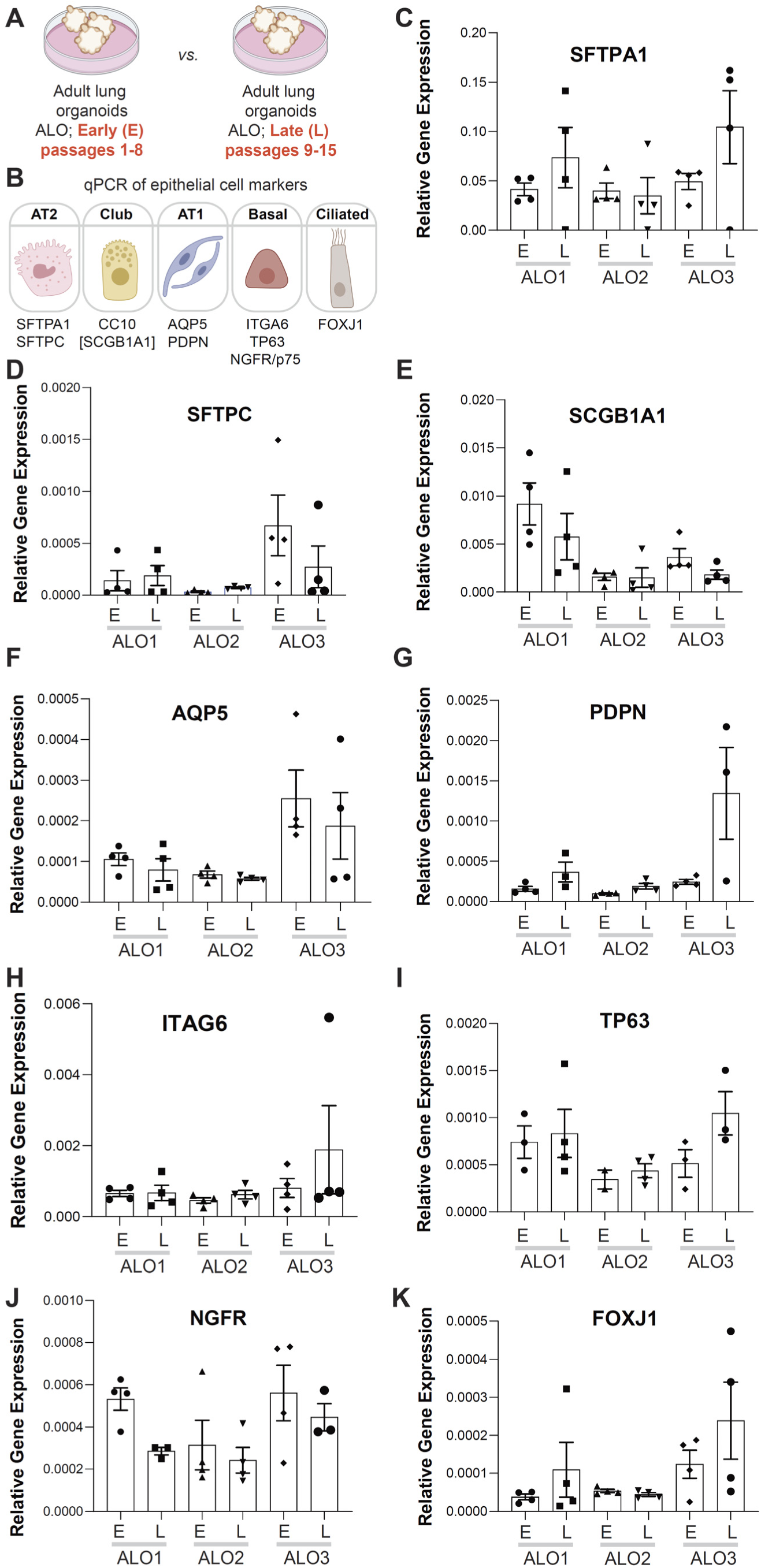
**Adult stem cell-derived lung organoids (ALO) generally maintain their cellular composition from early (E) to late (L) passages, as determined by cell type specific gene expression by qPCR.** **A-B.** Schematics depict the study goal in this figure, i.e., analysis of cell type specific transcripts in early (E) vs late (L) passages of ALO1-3 lines. **C-I.** Bar graphs display the relative abundance of various cell type markers (normalized to 18S) in adult lung organoids from either early (E) or late (L) passages of ALO lines 1-3. *p*-values were analyzed by one-way Anova. Error bars denote S.E.M; n = 3-6 datasets. Statistically significant differences were not noted in any of the transcripts analyzed.

**Figure 2- Figure Supplement 5.**
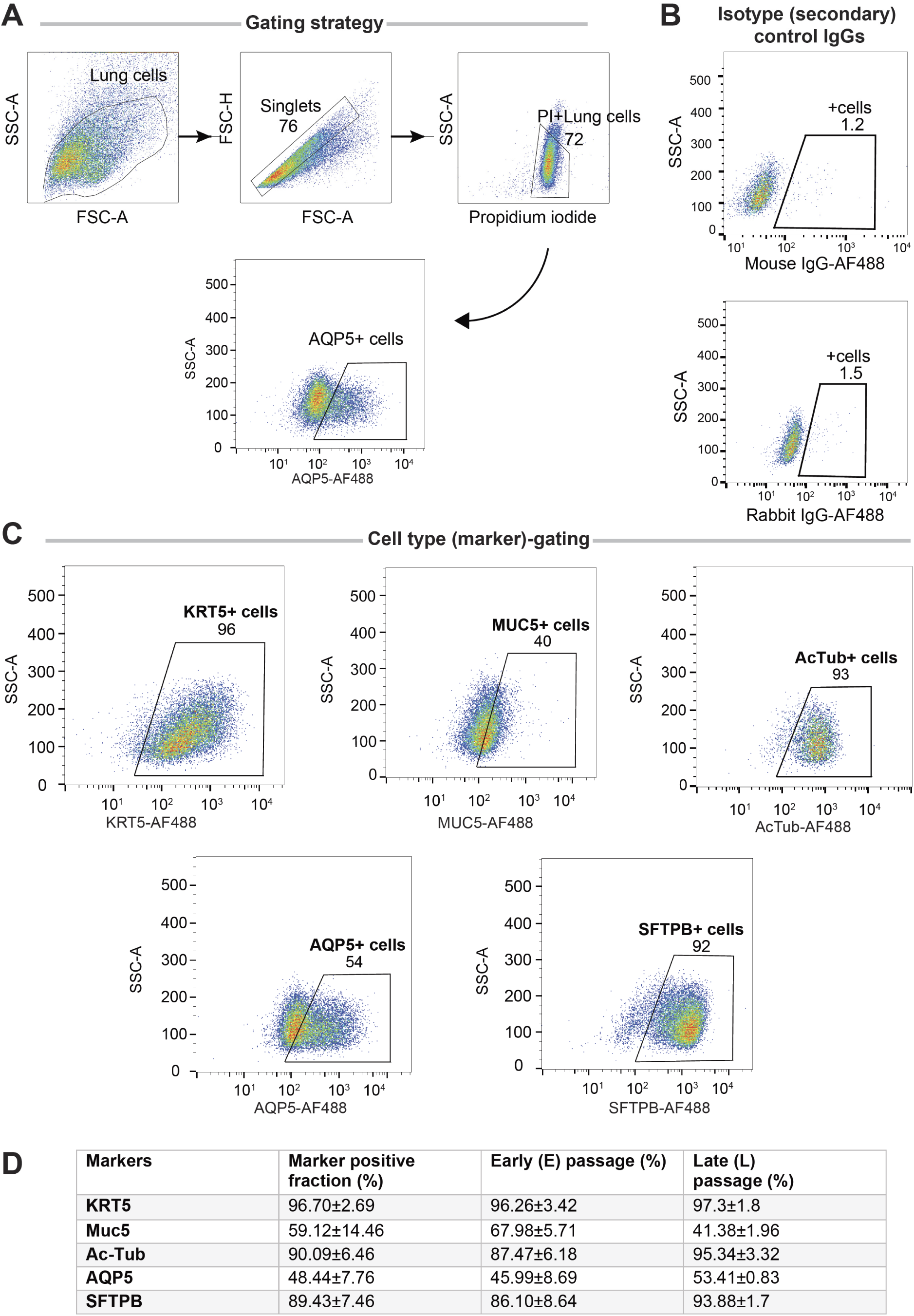
**Adult stem cell-derived lung organoids (ALO) are comprised of both proximal and distal airway epithelial population and generally maintain such diversity from early (E) to late (L) passages, as determined by FACS**. Lung monolayers were dissociated into single cells and analyzed using flow cytometry. Gating strategy depicted in A, isotype controls in B and panel C show various lung cell types. Numbers denote %. Table in D lists marker-positive cell fractions in ALO1-3, presented either as averaged over both early and late passages combined (column 2), or separated into early (column 3) or late (column 4) passages. These findings are consistent with others’ findings by multi-channel FACS^114^showing that although many of these markers are highly expressed in a certain cell type, they are shared at lower levels among other cell types.

**Figure 3- Figure Supplement 1.**
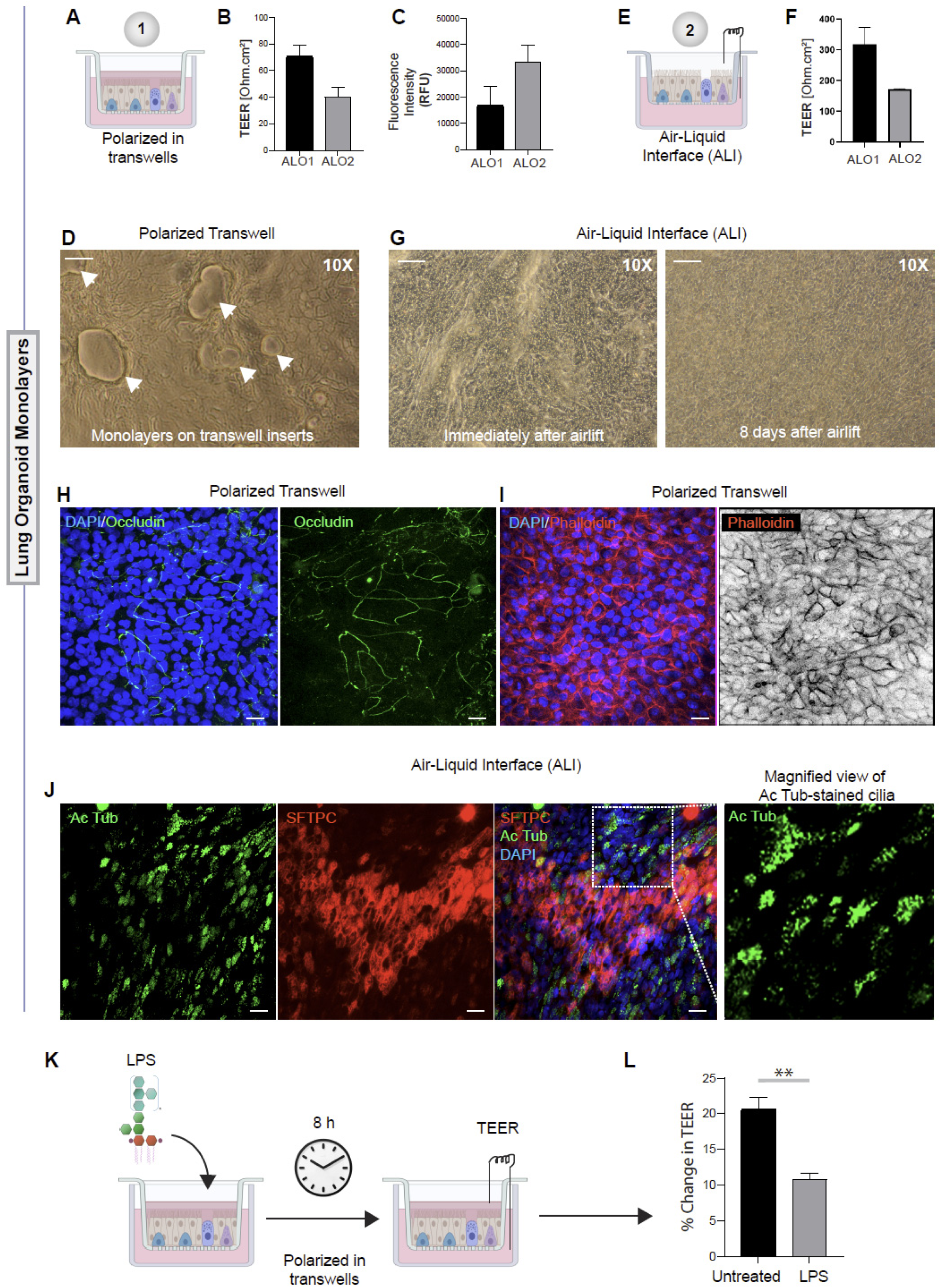
**Monolayers derived from adult lung organoids (ALO) can form an epithelial barrier.** **A-G**. Two different types of 2D polarized monolayers are prepared using adult lung organoids. Schematics in A and E show growth as submerged or ALI models, respectively. Panel B shows bar graphs with transepithelial resistance (TEER) across submerged monolayers grown in transwells. Panel C shows bar graphs for relative fluorescence unit (RFU) of the FITC-labeled dextran flux from the apical to basolateral chambers of a submerged monolayer. D. Brightfield images show representative fields of submerged monolayers grown on transwells. Scale bar = 100 µm. Arrows = self-organized vacuolar regions were seen. F. Bar graphs with transepithelial resistance (TEER) across ALO-derived monolayers grown as ALI models. G. Brightfield images show representative fields of ALI monolayers at two different time points during culture. Scale bar = 100 µm. **H-I**. Submerged monolayers of ALO were fixed with methanol (H) or paraformaldehyde (I) prior to co-staining with DAPI (blue; nuclei) and either occludin (green, H) or phalloidin (red; I). Scale bar = 20 µm. **J**. ALO monolayers were grown as ALI models were fixed and co-stained for SFTPC (red), Ac-Tub (green) and DAPI (blue; nuclei) and visualized by confocal immunofluorescence microscopy. Scale bar = 20 µm. **K-L**. Schematic in K shows the study design for challenging submerged monolayers with 500 ng/ml LPS, followed by TEER measurement. Bar graphs in L display the % change in TEER observed with or without LPS treatment normalized to the baseline TEER. *p*-values were analyzed by one-way Anova. Error bars denote S.E.M; n = 3-6 datasets. **p = <0.01.

**Figure 3- Figure Supplement 2.**
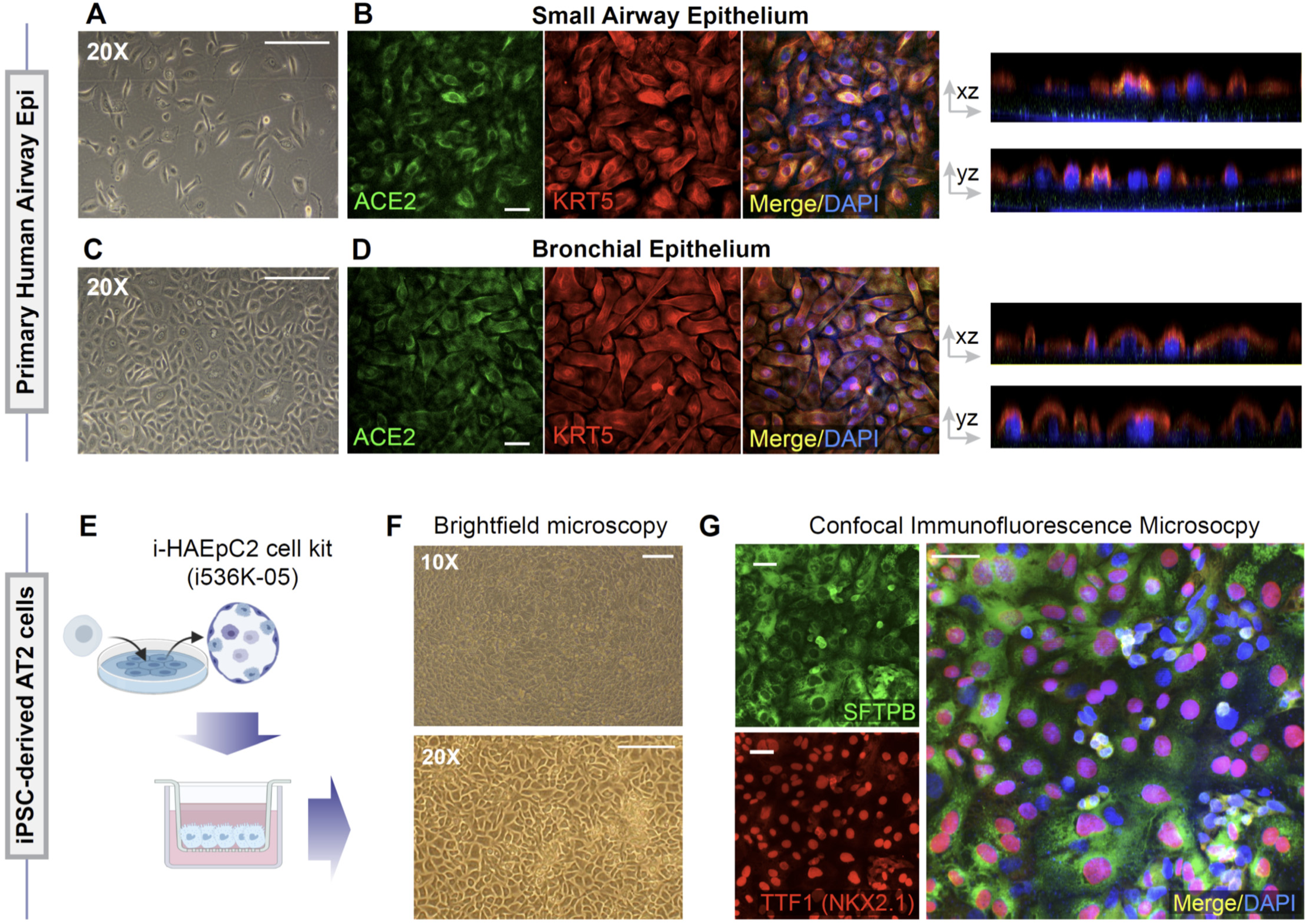
**Alternative models of lung epithelial cells used in this work for modeling SARS-CoV-2 infection and/or as a control for gene expression studies.** **A-D**. Monolayers of primary airway epithelial cells (small airway epi; A-B; bronchial epi; C-D) were visualized by bright field microscopy (A, C) or by fixing, staining and visualizing by confocal microscopy (B, D). Representative images in B and D are presented as maximum projected z-stacks on the left and as an orthogonal view on the right. **E-G**. hiPSC-derived AT2 cells, prepared using the i-HAEpC2 cell kit, were grown in monolayers on transwell inserts to form a polarized. Brightfield images are shown in F. Monolayers were fixed and stained for several markers and analyzed by confocal microscopy. Representative images are shown in G. Scale bar = 20 µm.

**Figure 3- Figure Supplement 3.**
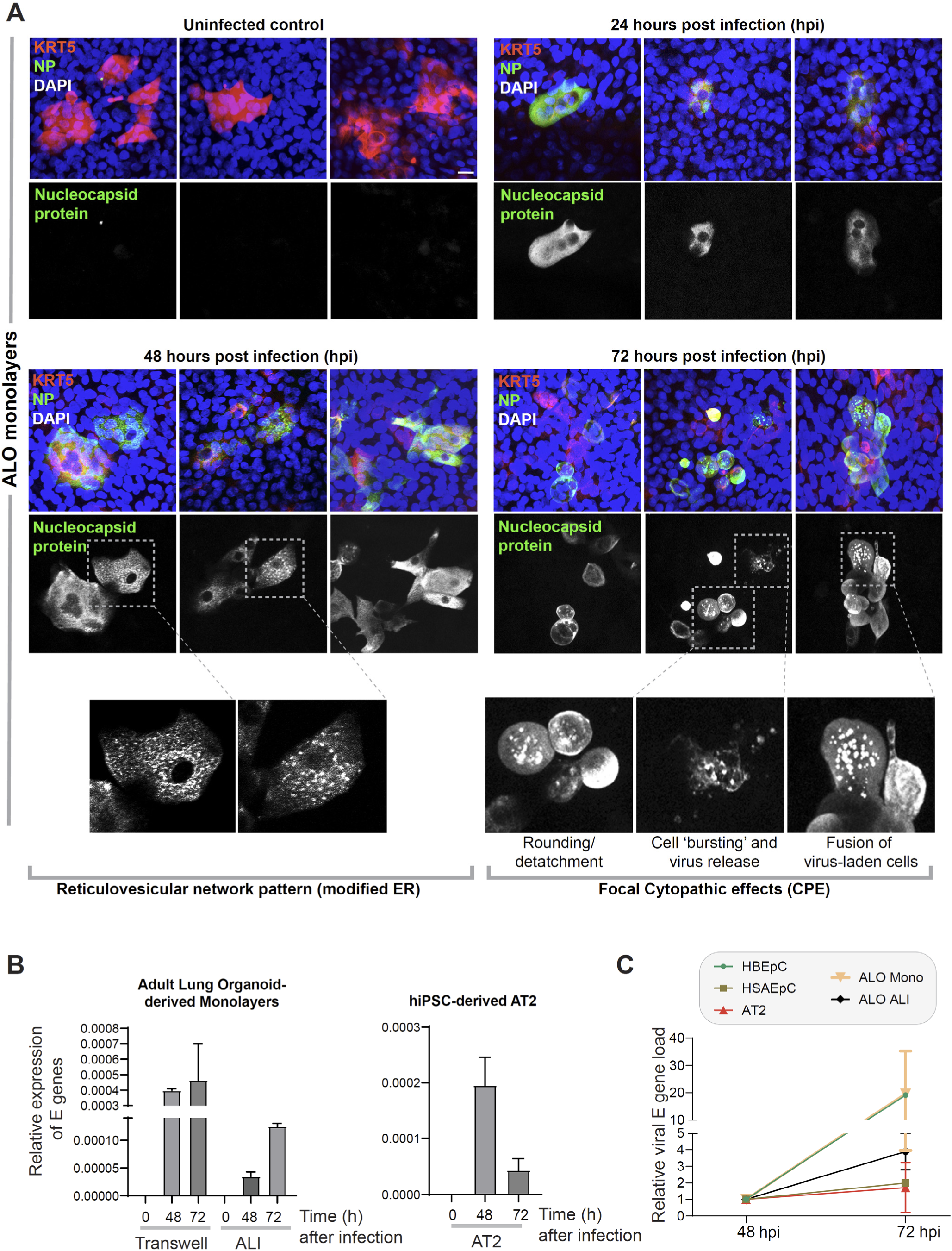
**Proof of SARS-CoV-2 infectivity.** **A.**Monolayers of ALO1-3 were challenged with SARS-CoV-2 for indicated time points prior to fixation and staining for KRT5 (red) and viral nucleocapsid protein (green) and DAPI (blue; nuclei) and visualized by confocal microscopy. Representative images are shown, displaying various cytopathic effects. Scale bar = 15 µm. **B.** Monolayers of ALO (either transwell submerged models or ALI, *Left*) and monolayers of hiPSC-derived AT2 cells (*Right*) were infected or not with SARS-CoV-2 and analyzed for viral envelope gene (E gene). Bar graphs display the relative expression of E gene in infected ALO monolayers, indicative of viral infection. **C.** Line graphs show the change in E gene expression in infected monolayers over 24 h period (from 48 hpi to 72 hpi) where values at 72 hpi are normalized to that at 48 hpi. Data is presented as S.E.M. of 3 independent repeats.

**Figure 4- Figure Supplement 1.**
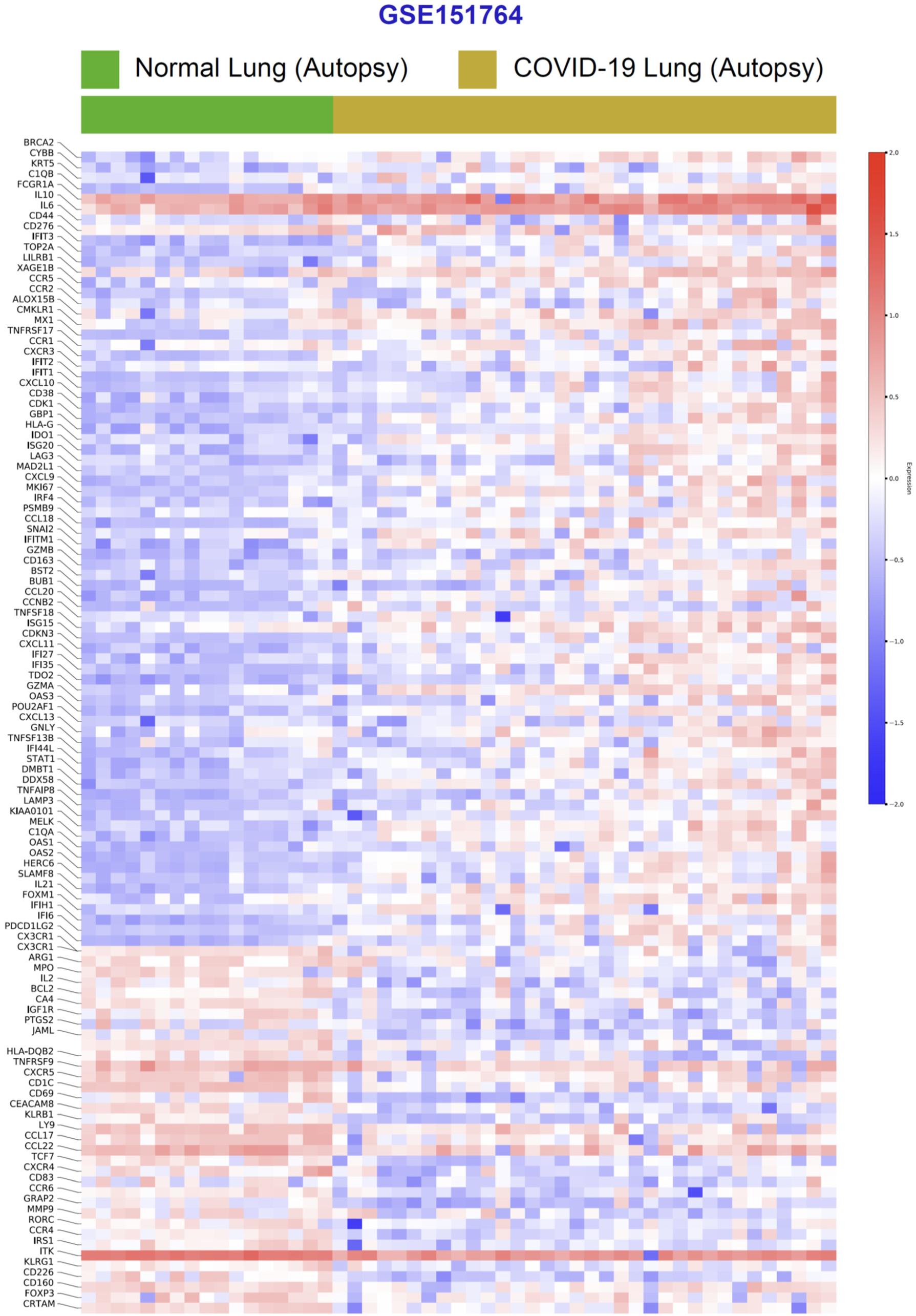
**Differential expression analysis of RNA Seq datasets from lung autopsies (normal vs. COVID-19).** Publicly available RNA Seq datasets (GSE151764) of lung autopsies from patients who were deceased due to COVID-19 or non-infectious causes (normal lung control) were analyzed for differential expression of genes and displayed as a heatmap.

**Figure 4- Figure Supplement 2.**
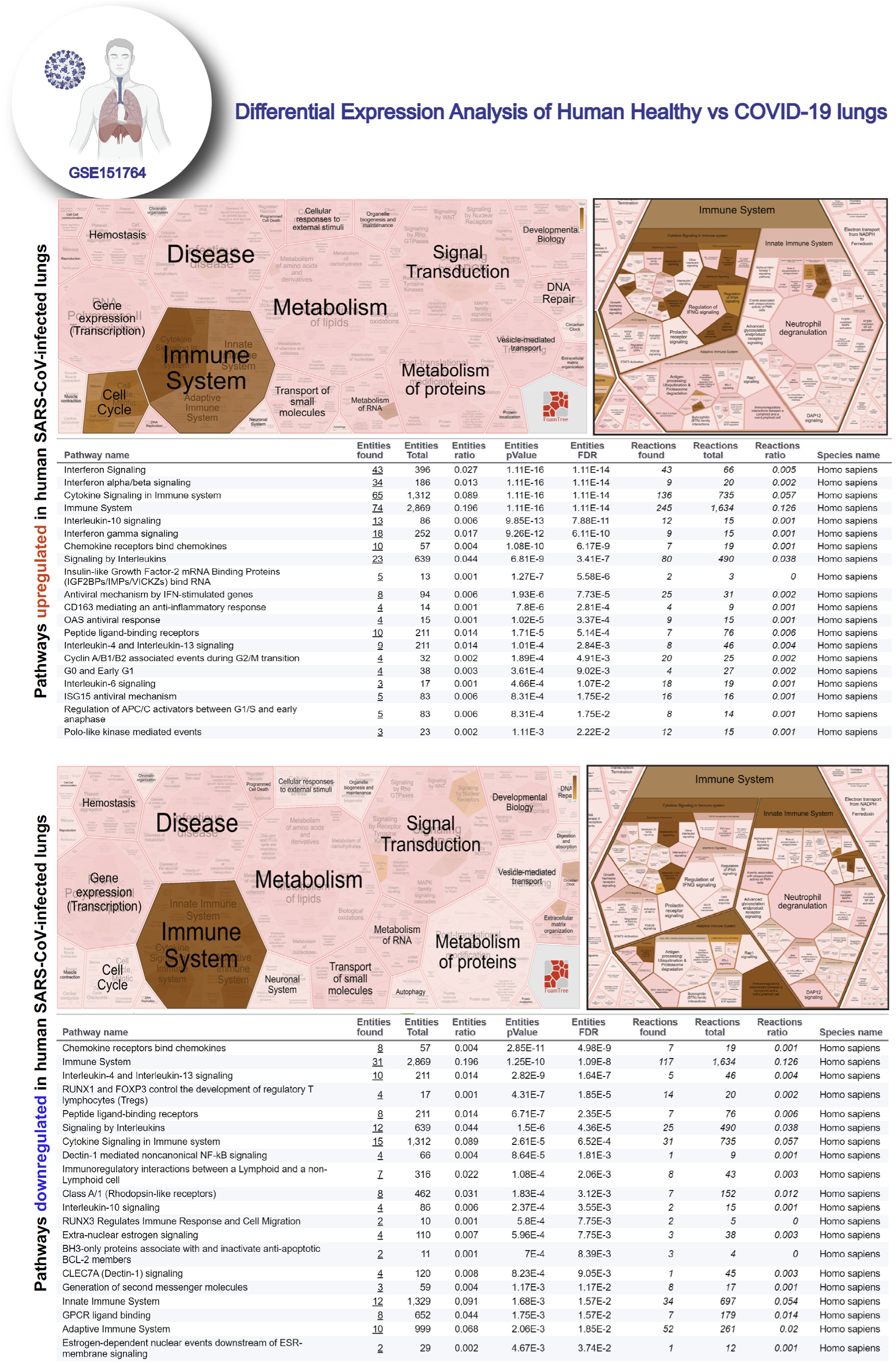
**Reactome pathway analysis of differentially expressed genes in lung autopsies (normal vs. COVID-19).** Reactome-pathway analysis of the differentially expressed genes shows the major pathways upregulated in COVID-19-affected lungs. *<u>Top</u>*: Visualization as flattened (left) and hierarchical (right, insets) reactome. *<u>Bottom</u>*: Visualization of the same data as tables with statistical analysis indicative of the degree of pathway enrichment.

**Figure 5- Figure Supplement 1.**
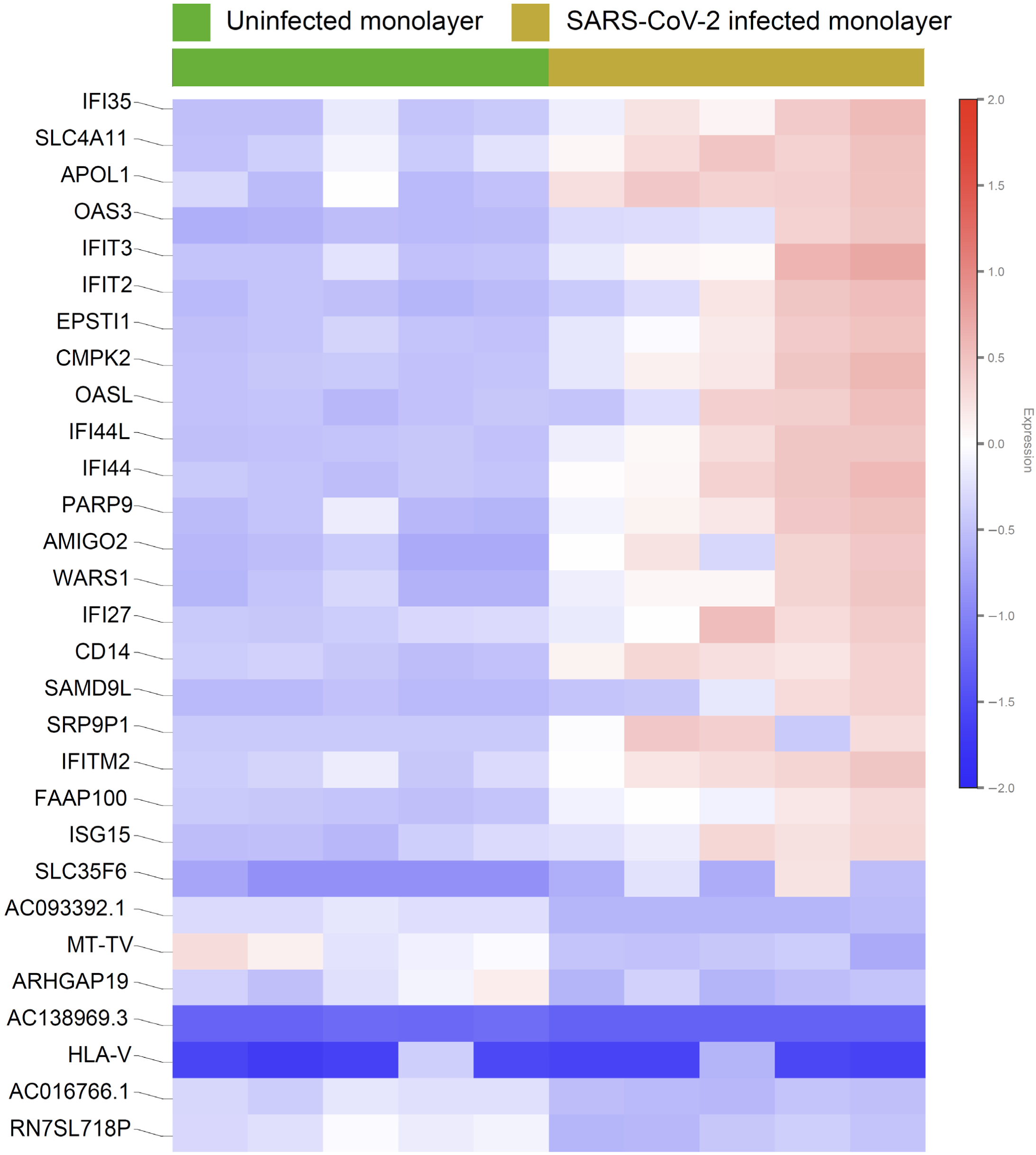
**Differential expression analysis of RNA Seq datasets from adult lung organoid monolayers, infected or not, with SARS-CoV-2.** Adult lung organoid (ALO)-derived grown in transwells as submerged monolayers were infected or not with SARS-CoV-2 were analyzed by RNA seq and differential expression analysis. Differentially expressed genes are displayed as a heatmap.

**Figure 5- Figure Supplement 2.**
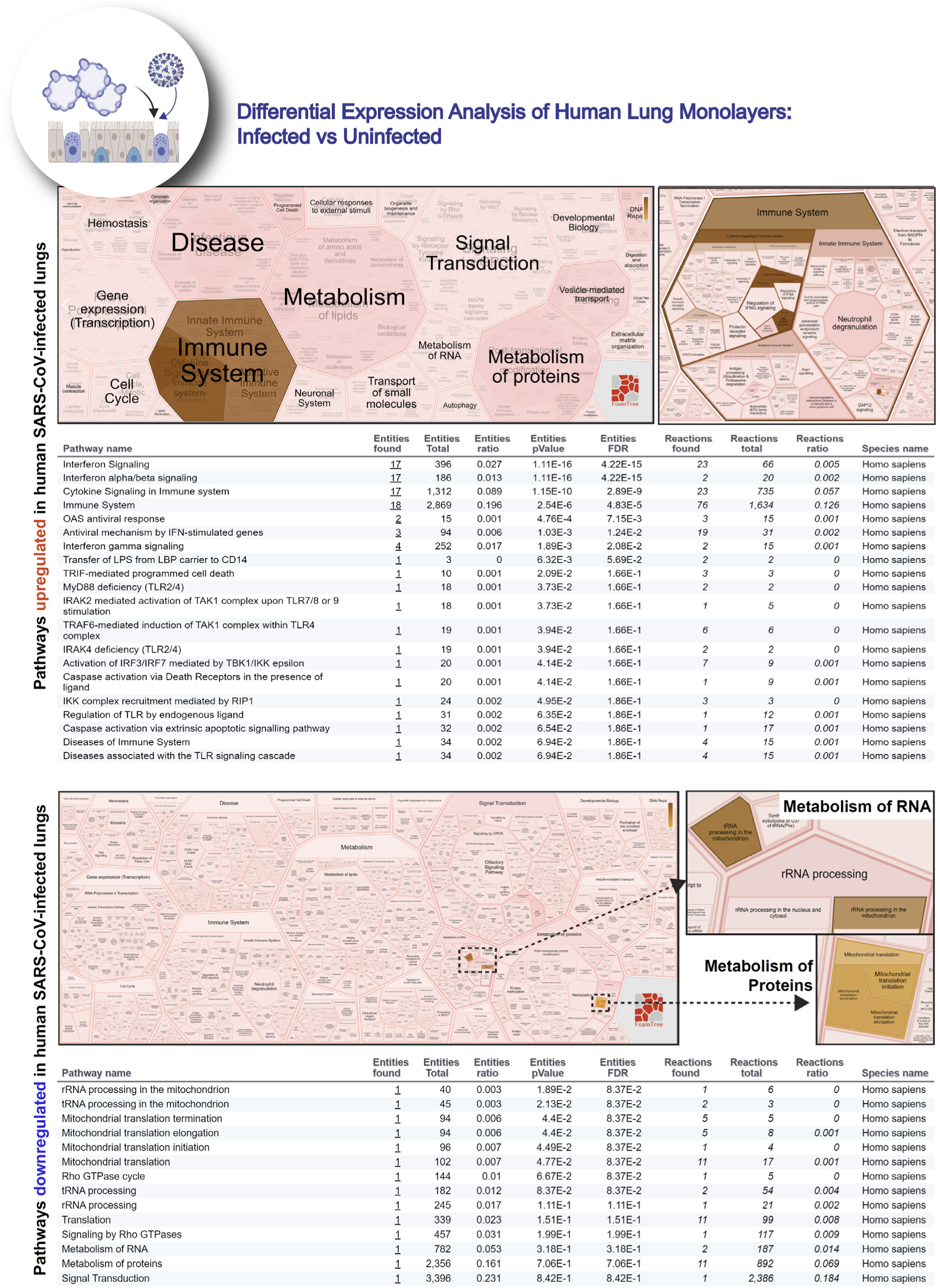
**Reactome pathway analysis of differentially expressed genes in lung organoid monolayers infected with SARS-CoV-2.** Reactome-pathway analysis of the differentially expressed genes shows the major pathways upregulated in SARS-CoV-2-infected lung organoid monolayers. *<u>Top</u>*: Visualization as flattened (left) and hierarchical (right, insets) reactfoam. *<u>Bottom</u>*: Visualization of the same data as tables with statistical analysis indicative of the degree of pathway enrichment.

**Figure 5- Figure Supplement 3.**
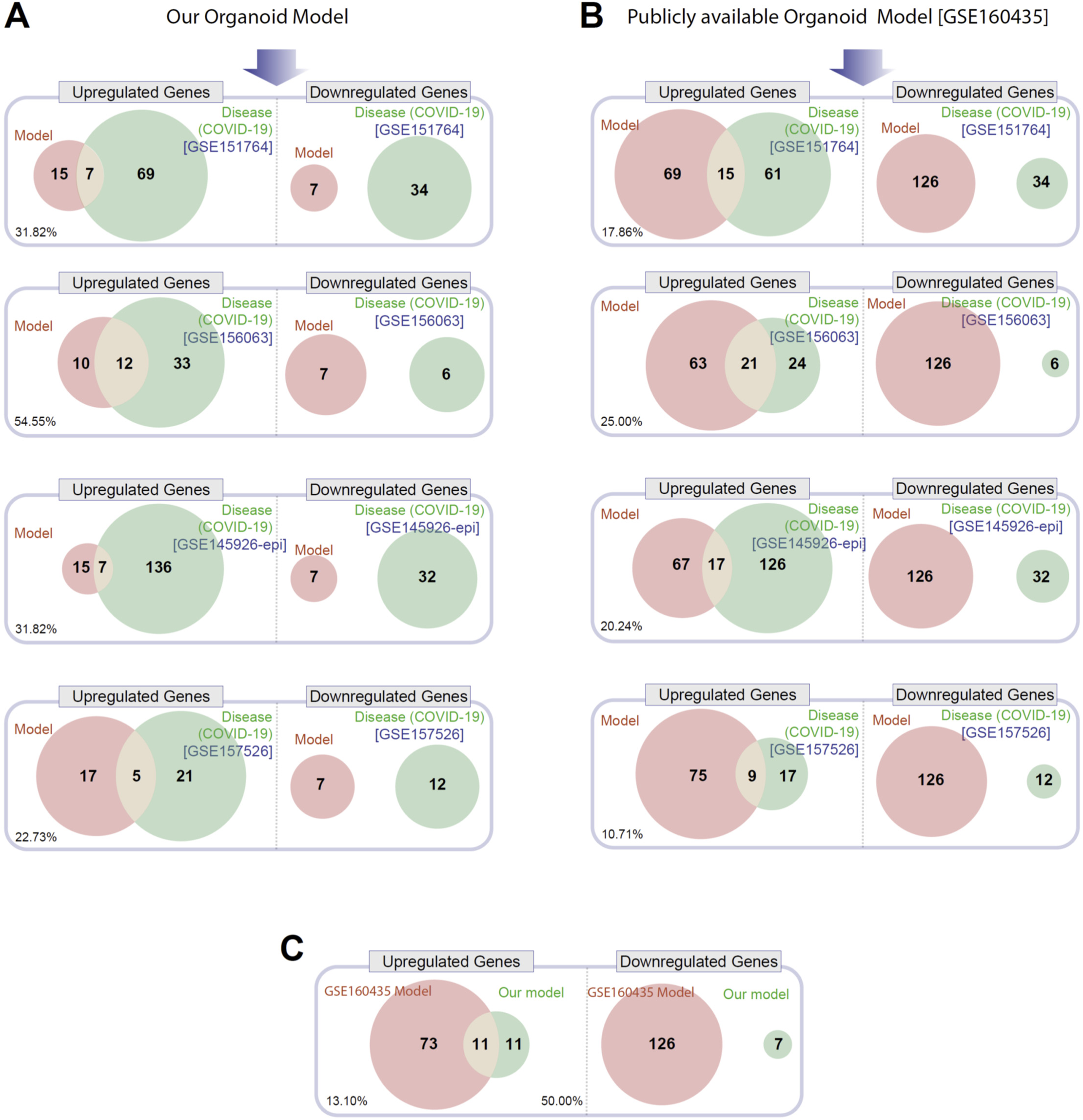
**Head-to-head comparison of our adult lung organoid (ALO)-derived model of COVID-19 versus another lung organoid model in their ability to recapitulate the DEGs observed in lung tissues from fatal cases of COVID-19.** **A.**Venn diagrams show the number of overlapping and non-overlapping differentially expressed genes (DEGs, both Up and Downregulated genes) between our organoid model and four human COVID-19 patient derived samples (left). GSE151764 represents postmortem COVID-19 and normal lung tissues; GSE156063 represents upper airway samples from patients with COVID-19; GSE145926 represents sorted epithelial population from bronchoalveolar lavage fluid (BALF) derived from patients with varying severity of COVID-19; GSE157526 represents tracheal-bronchial cells infected with SARS-Cov2. **B**. Venn diagrams as in A, comparing a publicly available SARS-Cov2 infected human lung organoid model (GSE160435) and the same four human COVID-19 respiratory cohorts as in A. **C**. Venn diagrams show the DEGs between our organoid model and the publicly available lung organoid model. The comparison was carried out by calculating the percentage of the common Up/Down DEGs represented within the total Up/Down DEG for the two models in each Venn diagram.

